# Identification of novel myokines and putative protein targets that mediate functional adaptations in response to chronic contractile activity induced skeletal muscle-extracellular vesicle treatment

**DOI:** 10.1101/2024.09.09.612156

**Authors:** Patience O. Obi, Tamiris F. G. Souza, Ying Lao, Kirk J. McManus, Joseph W. Gordon, Richard LeDuc, René P. Zahedi, Ayesha Saleem

## Abstract

We have previously shown that skeletal muscle-derived extracellular vesicles (EVs) released post-chronic contractile activity (CCA) increased mitochondrial biogenesis in murine myoblasts, and decreased cell viability and induced apoptosis and senescence in non-small cell lung cancer cells. While the underlying mechanisms are unknown, the effects perpetuated were dependent on membrane-bound proteins. Here, we performed an extensive LC-MS/MS proteomic analysis on EVs from control and CCA myotubes. A total of 2900 proteins were identified in CON-EVs and CCA-EVs, including EV-associated proteins such as TSG101, tetraspanins (CD9, CD81, and CD63), flotillin-1, and annexins. Of these, 856 proteins are novel and not listed in EV databases (ExoCarta and Vesiclepedia), indicating that myotube-EVs harbor proteins not yet identified in EVs of different origin. Additionally, we identified 2062 unique proteins that have not yet been previously reported in myotube-EVs to date. Remarkably, of the 2900 total proteins identified, we observed 46 upregulated, and 25 downregulated differentially expressed proteins (DEPs) in CCA-EVs *vs*. control-EVs. Most of upregulated DEPs include EV-associated proteins. Comparing the 71 DEPs with proteins expressed in skeletal muscle indicated 61 of these as potential myokines. We identified actin cytoskeleton signaling, integrin signaling and muscle contraction as the most enriched pathways among the DEPs using different databases/software including FunRich, KEGG, STRING and Ingenuity Pathway Analysis. Using a relevance score that prioritized membrane-bound proteins with known function in mitochondrial biogenesis and inhibition of cancer growth, we identified top-scoring highly enriched DEPs of interest: IGF1R, ATP7A, PFN1, GJA1, PRKCA and ITGA6. We confirmed upregulation of these targets in EVs using immunoblotting. Among these top-scoring DEPs, PFN1, and ITGA6 are associated with EVs, with expression upregulated following acute exercise. In summary, we report the first comprehensive analysis of skeletal muscle-EV proteome following CCA, with identification of putative protein targets and signaling pathways that may execute the pro-metabolic and anti-tumorigenic effects of CCA-EVs.

## Introduction

Regular exercise is an important physiological stressor, conferring numerous systemic benefits, including enhanced mitochondrial function, protection against cardiovascular diseases, and potential anti-tumorigenic effects, among many other beneficial adaptations [1]–[3]. Several organ systems including skeletal muscle [4], adipose tissue [5], brain [6], bone [7], and liver [8] secrete proteins in response to exercise, that may contribute to these systemic benefits. However, skeletal muscle is the primary organ responsible for regulating the majority of exercise-induced adaptations [9]. As the largest organ system in the body comprising 40% of body mass, skeletal muscle plays a crucial role in regulating whole-body metabolic capacity. Substantial evidence suggests that the systemic adaptative effects of exercise are partly mediated by the release of myokines from skeletal muscle during exercise [10], [11]. Myokines, which are typically proteins, exert their effects on various organs by acting in an autocrine, paracrine, or endocrine fashion. Examples include interleukin (IL)-6, IL-10, IL-15, irisin, meteorin, myostatin, among others [11]. In addition to myokines, skeletal muscle can also release extracellular vesicles (EVs) during exercise.

Extracellular vesicles (EVs) are integral to cell-cell communication and are present in all biological fluids, including blood, urine, cerebrospinal fluid, saliva, breast milk, and amniotic fluid, among others [12], [13]. Traditionally, EVs are categorized into three subtypes based on their biogenesis, size, and biological properties. They include exosomes (30-150 nm), which are formed through the invagination of endosomal membranes resulting in the formation of intraluminal vesicles within multivesicular bodies (MVBs) that are released via exocytosis; microvesicles (100-1000 nm), which originate from the outward budding of the plasma membrane; and apoptotic bodies (500-5000 nm), which are generated from the outward blebbing of an apoptotic cell membrane [14]. According to the Minimal Information for Studies of EVs (MISEV) 2023 guidelines, EVs should be classified based on: i) size, where small EVs (sEVs) are <200 nm and large EVs (lEVs) >200 nm; ii) biochemical composition; and, iii) cell of origin [15]. EVs carry a variety of biological molecules, including proteins, DNA, mRNA, miRNA, lipids, and metabolites, facilitating the transfer of this cargo from donor to recipient cells [16]. Over the past few decades, the role of EVs in mediating intercellular communication under both physiological and pathological conditions, such as exercise, cancer, and metabolic diseases, has been a significant focus of research.

Previous studies have shown that acute/chronic exercise in humans, mice and rats increases the release of EVs in the circulation [17]–[26]. To fully explore the functional effects of exercise derived EVs in health and disease, it is important to measure the cargo content of the EVs, and delineate their functional role in downstream effects. We have previously shown that chronic contractile activity (CCA) increased secretion of Skm-EVs from stimulated myotubes, and these in turn increased mitochondrial biogenesis in murine myocytes [27], and reduced cell viability, induced apoptosis and senescence in non-small cell lung cancer cells [28]. Notably, we also demonstrated that these effects were abrogated when CCA-EVs were pre-treated with Triton X-100 (Triton) and proteinase K (Pro K), indicating that the effects were likely mediated by the membrane-bound (transmembrane or peripheral membrane-associated) EV proteins and/or EV corona. Based on these observations, we sought to examine the proteome of Skm-EVs released post-CCA to identify specific EV proteins that could explain the observed pro-metabolic, anti-tumorigenic effect of CCA-EV treatment. Here, we report findings from the first comprehensive, quantitative proteomic analysis on EVs derived from control and CCA exposed myotubes. We observed a CCA-induced increase in several classes of proteins associated with small EVs. We further validated the upregulation of several differentially expressed proteins associated with mitochondrial biogenesis and/or inhibition of cancer growth by Western blotting. Our study offers novel insights into the proteome of Skm-EVs derived after CCA, and identifies several potential protein candidates that may be crucial for mediating the pro-metabolic and anti-tumorigenic effects of chronic endurance exercise as transmitted by EVs.

## Methods

### Cell culture

Murine C2C12 myoblasts were seeded in a six well plate pre-coated with 0.2% gelatin, and grown in fresh Dulbecco’s Modification of Eagle’s Medium (DMEM; Sigma-Aldrich) supplemented with 10% fetal bovine serum (FBS; Gibco/ThermoFisher Scientific) and 1% penicillin/streptomycin (P/S) (growth media). Cells were grown at 37 °C in 5% CO_2_ incubator for 24 h. When myoblasts reached approximately 90-95% confluency, the growth media was switched to differentiation media (DMEM supplemented with 5% heat-inactivated horse serum (HI-HS; Gibco/ThermoFisher Scientific) and 1% P/S) for 5 days to differentiate myoblasts into myotubes.

### Chronic contractile activity (CCA) in myotubes

CCA was performed using electrical pulse stimulation as previously described [27]. After five days of differentiation, myotubes were divided into control (CON) and chronic contractile activity (CCA) groups. The CCA plate was stimulated using the C-Pace EM Cell culture stimulator, with C-dish and carbon electrodes (IonOptix, Milton, MA, United States) while the CON plate had the C-dish placed on top but without the carbon electrodes. The contractile activity protocol was performed at a frequency of 1 Hz (at 2 ms pulses) and an intensity of 14 V chronically for 3 h/day for four consecutive days. After the first bout of contractile activity, spent media was changed to exosome-depleted differentiation media (DMEM supplemented with 5% exosome-depleted HI-HS and 1% P/S) for both CON and CCA groups and myotubes left to recover for 21 h. After recovery, the conditioned media was collected from the cells and used for EV isolation, after which they were stored in the -80 °C freezer. This process was repeated after each bout of contractile activity for 4 days. After the four day CCA protocol was completed, the EVs from days 1 – 4 were pooled together and used for mass spectrometry.

### Isolation of EVs by differential centrifugation

EV isolation by differential ultracentrifugation (dUC) was performed as previously described [27]. Briefly, conditioned media (CM) from each six well plate (12 mL) was collected and centrifuged at 300*xg* for 10 min at 4 °C to pellet dead cells (Sorvall™ RC 6 Plus Centrifuge, F13-14 fixed angle rotor), followed by centrifugation at 2000*xg* for 10 min at 4 °C to remove cell debris. The resulting supernatant was centrifuged at 10,000*xg* for 30 min at 4 °C to remove large vesicles. Using an ultracentrifuge (Sorvall™ MTX 150 Micro-Ultracentrifuge, S58-A fixed angle rotor), the supernatant was spun at 100,000*xg* for 70 min at 4 °C to obtain the exosome/sEV pellet. The exosome/sEV pellet was resuspended in 1 mL PBS and centrifuged again at 100,000*xg* for 70 min at 4 °C. After centrifugation, the final exosome/sEV pellet was resuspended in 50 µL PBS and used for subsequent analysis.

### Liquid chromatography (LC)-MS/MS analysis

Proteomic analysis of myotube-EVs was performed using a tandem mass tags (TMT)-based quantitative approach in collaboration with the Manitoba Centre for Proteomics and Systems Biology. Five distinct experiments, i.e., biological replicates were performed for both CON-EVs and CCA-EVs, and LC-MS/MS performed for each. To extract proteins, EVs were resuspended in lysis buffer (100 mM HEPES (pH 8.5), 3% SDS, 1X protease inhibitor cocktail). EVs were sonicated in a water bath three times for 15s per cycle with a 1-minute cooling period between each cycle. Protein concentration was determined using Pierce detergent compatible Bradford assay kit (Thermo Fisher Scientific) as before [29]. All protein samples were processed and handled using single-pot solid-phase-enhanced sample preparation (SP3) protocol. Prior to SP3 treatment, two types of carboxylate-modified SeraMag Speed beads (GE Life Sciences) were combined in a ratio of 1:1 (v/v), rinsed, and reconstituted in water at a concentration of 20 µg solids/µL. Initially, 25 µg of lysate was reduced with 10 mM (final concentration) DTT for 30 min at 60 °C followed by alkylation using 50 mM (final concentration) IAA for 45 min in dark at room temperature. After that, 3 µL of the prepared bead mix was added to the lysate and samples were adjusted to pH 7 using HEPES buffer. To promote proteins binding to the beads, acetonitrile was added to a final concentration of 70% (v/v) and samples were incubated at room temperature on a tube rotator for 18 min. Subsequently, beads were immobilized on a magnetic rack for 1 min. The supernatant was discarded, and the pellet was rinsed 2X with 200 µL of 70% ethanol and 1X with 200 µl of 100% acetonitrile while on the magnetic rack. Rinsed beads were resuspended in 15 µL of 50 mM HEPES buffer (pH 8) supplemented with trypsin/LysC mix (Promega) at an enzyme-to-protein ratio of 1:25 (w/w) and incubated for 16 h at 37 °C. After overnight digestion, peptides were eluted and immediately labeled with TMT. TMT labeling was performed as specified by the manufacturer (Thermo Scientific), except that TMT were dissolved in DMSO. Peptide concentration was determined using Pierce Quantitative Fluorometric Peptide Assay (Thermo Fisher Scientific). Equivalent labeled samples within each TMT set were mixed prior to 1D LC-MS/MS. Agilent 1100 series LC system with UV detector (214 nm) and 1mm×100mm XTerra C18, 5 µm column (Waters, Ireland) was used for pH 10 first dimension reversed-phase separation. Both eluents A (water) and B (1:9 water: acetonitrile) contained 20 mM ammonium formate pH 10. 1.80% acetonitrile per min gradient (0.1-59.9% acetonitrile in 30 min) was delivered at flow rate of 150 µL/min. Twenty 1-min fractions were collected and concatenated into 10 to provide optimal separation orthogonality. These fractions were lyophilized and resuspended in 0.1% formic acid for the second dimension. Analysis of TMT labeled peptide digests was carried out on an Orbitrap Exploris 480 instrument (Thermo Fisher Scientific, Bremen, Germany). The individual fractions were introduced using an Easy-nLC 1000 system (Thermo Fisher Scientific) at 1.5 µg per injection. Mobile phase A was 0.1% (v/v) formic acid and mobile phase B was 0.1% (v/v) formic acid in 80% acetonitrile (LC-MS grade). Gradient separation of peptides was performed on a C18 (Luna C18(2), 3 µm particle size (Phenomenex, Torrance, CA)) column packed in-house in Pico-Frit (100 µm X 30 cm) capillaries (New Objective, Woburn, MA). Peptide separation was using the following gradient: started with 3% of phase B, 3-7% over 5 min, 7-25% over 59 min, 25-60 % over 10 min, 60-90% over 1 min, with final elution of 90% B for 15 min at a flow rate of 300 nL/min. Data acquisition on the Orbitrap Exploris 480 instrument was configured for data-dependent method and operated in a positive mode with the FAIMS Pro interface. The inner and outer electrodes of FAIMS were heated to a common temperature of 100 °C to maximize ion transmission. A standard nitrogen flow of 4.1 L/min is used for all experiments with user gas kept at 0 L/min. Parallel experiments were created for CV -50 and -70, with 1.7s and 1.3s cycle time for sequential survey scans and MS/MS cycles, respectively. Spray voltage was set to 1.9 kV, funnel RF level at 40, and heated capillary at 275 °C. Survey scans covering the mass range of 375– 1500 m/z were acquired at a resolution of 120,000 (at m/z 200), with a maximum ion injection time of 50 milliseconds, and a normalized automatic gain control (AGC) target of 200%. This was followed by MS2 acquisition at a resolution of 45,000, selected ions were fragmented at 36% normalized collision energy, with intensity threshold kept at 1e4. AGC target value for fragment spectra was set to 300%, with a maximum ion injection time set to auto and an isolation width set at 0.7 m/z. Precursor Fit is turned on with fit threshold and fit window set to 70% and 0.7, respectively. Dynamic exclusion of previously selected masses was enabled for 25s, charge state filtering was limited to 2–5, peptide match was set to preferred, and isotope exclusion was on.

### Data processing

Raw data was analyzed using Proteome Discoverer v2.5 (Thermo Scientific). Sequest HT was used for database searching against a murine Swissprot database (downloaded July 18, 2022, 17,096 forward sequences) and a database with common contaminations, using the following settings. MS and MS/MS tolerances were 10 ppm and 0.02 Da, trypsin was used as enzyme with a maximum of 2 missed cleavages. Oxidation of Met (+15.995 Da), deamidation of Asn, Gln (+0.984 Da) were used as variable modifications. Carbamidomethylation of Cys (+57.021 Da), and TMT-labeling (+229.163 Da) of N-termini and Lys residues were used as fixed modifications. The reporter ion node was used for TMT quantitation, percolator was used to calculate posterior error probabilities and all data was filtered to <1% false discovery rate (FDR) on peptide and protein levels. Only proteins that were quantified with at least 1 protein unique peptide were kept. Perseus v2.0 was used to log2-transform protein reporter ion intensities. Statistically significantly differential proteins were determined at 5% FDR after multiple testing correction, as indicated in the Volcano plot.

### Bioinformatics analysis

The volcano plot and heatmaps were generated using Perseus v2.0 and GraphPad prism v10.2.2 to demonstrate the expression of proteins. Functional enrichment analysis for gene ontology (GO) annotations and comparison of data sets was conducted using FunRich (http://www.funrich.org/, v3.1.4, accessed on April 12, 2024) [30]. Since FunRich is a human database, the rodent proteome was downloaded from UniProt. Comparisons between our EV proteome and EV databases (ExoCarta and Vesiclepedia) were conducted by importing the online protein data from these databases (as recent as April 17, 2024) into FunRich. This data set included all proteins listed in ExoCarta (http://www.exocarta.org/) [31] and Vesiclepedia (http://microvesicles.org/) [32]. GO analysis of annotated proteins was performed for cellular component, biological process and molecular function. Enriched GO terms that were significant (GO terms with a p value <0.05) in FunRich were selected. Functional analysis was also conducted based on KEGG analysis (http://bioinformatics.sdstate.edu/go/) and the top enriched pathways were based on enrichment. The interactions of differentially expressed proteins (DEPs) were evaluated using search tool for the retrieval of interacting genes and proteins (STRING) database (https://string-db.org/, version 12.0, accessed on April 17, 2024) [33]. The following settings were used for the STRING database. Organism: *Mus musculus*; network type: full STRING network; minimum required interaction score: medium confidence (0.400); max number of interactors to show: none; FDR stringency: medium (5 percent). Additionally, DEPs were uploaded into Qiagen’s Ingenuity pathway analysis (IPA) system to identify protein networks, canonical pathways, and diseases and functions that are most significant [34].

### Pro K-Triton EV treatment

To identify the location to CCA-EV proteomic cargo, CCA-EVs were treated with or without 0.1% Triton at room temperature, and incubated with 10 µg/mL Pro K (Sigma Aldrich) for 1 h at 37 °C as detailed before [27]. Pro K activity was inhibited by adding 5 mM phenylmethylsulfonyl fluoride (PMSF; Sigma Aldrich) for 10 min at room temperature. After, samples were washed with 1 mL filtered 1XPBS and loaded onto the Ultra-4 Centrifugal Filter columns (Amicon), and centrifuged at 14,000*xg* for 1 h at 4 °C, until ∼40 µL of concentrated sample remained. Concentrated EV fraction was used for protein extraction and western blotting as described below.

### Western blotting

EVs were lysed using 1:1 Pierce RIPA solution with protease inhibitor tablet (Roche) and EV concentration was determined using Pierce™ MicroBCA protein assay kit (ThermoFisher Scientific) as previously described [27]. Total protein (3-5 µg) from EV lysates were resolved on a 12-15% SDS-PAGE gel and subsequently transferred to nitrocellulose membranes. Membranes were blocked for 1h with 5% skim milk in 1× Tris-buffered saline-Tween 20 solution (TBST) at room temperature, followed by incubation with primary antibodies in 1% skim milk overnight at 4 °C. The following primary antibodies were used: rabbit anti-Connexin 43 or GJA1 (3512, Cell Signaling, 1:200), mouse anti-ATP7A (S60-4, Novus Biologicals, 1:200), rabbit anti-ITGA6 (3750, Cell Signaling, 1:200), rabbit anti PKCα or PRKCA (2056, Cell Signaling, 1:200), rabbit anti-IGF-I Receptor β (3027, Cell Signaling, 1:200), and rabbit anti-Profilin-1 (3237, Cell Signaling, 1:200). Subsequently, membranes were washed three times for 5 min with TBST, followed by incubation with anti-mouse (A16017, ThermoFisher) or anti-rabbit (A16035, ThermoFisher) IgG horseradish peroxidase secondary antibody (1:10,000) in 1% skim milk for 1h at room temperature, after which the membrane is washed again three times for 5 min each with TBST. Membranes were developed using enhanced chemiluminescence detection reagent (Bio-Rad Laboratories), and the films were scanned using the ChemiDoc^TM^ MP Imaging System (Bio-Rad Laboratories). Band intensities were quantified using the Image Lab Software (Bio-Rad Laboratories) and corrected for loading using ponceau S staining or coomassie blue.

### Statistical analysis

All data were analyzed using unpaired Student’s t-test, and individual data points are plotted, with mean ± standard error of mean (SEM) shown as applicable. All graphs were created using GraphPad-Prism software (version 10.2.2, GraphPad, San Diego, CA, USA). Significance was set at p ≤ 0.05. Mass spectrometry was performed on five biological replicates of CON-EV and CCA-EV, while the western blot analysis was performed on four biological replicates.

## Results

### Quantitative analysis of differentially expressed proteins in CCA-EVs vs. CON-EVs

A brief schematic workflow for the quantitative proteomics is shown in **Fig. 1**. In-depth proteomic analysis was performed to determine the proteome of CON-EVs and CCA-EVs. A total of 2900 proteins were identified and unsupervised hierarchical clustering illustrated as a heatmap (**Fig. 2A**) The full list of proteins is provided separately (**Table S1**). Differentially expressed proteins (DEPs) between CON-EVs and CCA-EVs were quantified and illustrated in a volcano plot, along with the non-statistically significant proteins (**Fig. 2B**). While overall proteome patterns between CON-EVs and CCA-EVs were similar, we identified a total of 71 proteins with an FDR<0.05 that were significantly different between CCA-EVs vs. CON-EVs (full list of proteins in **Table 1**). These proteins enabled a distinct separation of CON-EVs and CCA-EVs through unsupervised clustering (**Fig. 3A**). Among the DEPs, 46 proteins were significantly upregulated, and 25 proteins were significantly downregulated in CCA-EVs vs. CON-EVs.

**Figure 1.**
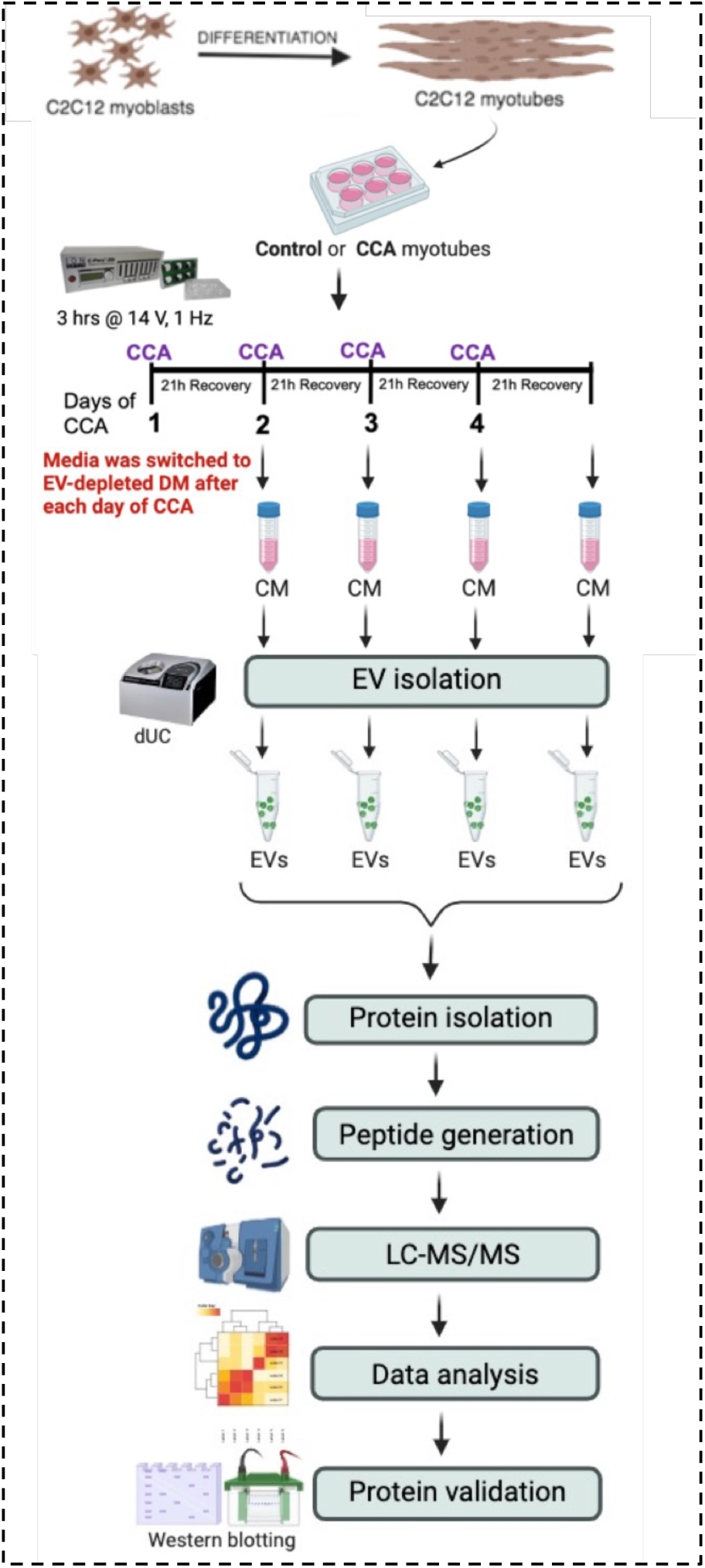
Overall study design. C2C12 myoblasts were fully differentiated into myotubes (MTs). MTs were divided into control (CON) and chronic contractile activity (CCA) plates. CCA-MTs were electrically paced for 3h/day x 4 days at 14 V to mimic chronic endurance exercise *in vitro* using IonOptix ECM. After each bout of contractile activity, we switched media to exosome-depleted differentiation media in both CON-MT and CCA-MT, and cells were allowed to recover for 21 h. EVs were isolated using dUC from conditioned media (CM) from control or stimulated myotubes. CON-EVs or CCA-EVs isolated after each bout of contractile activity were pooled together and analyzed by LC-MS/MS. Created with BioRender.com.

**Figure 2.**
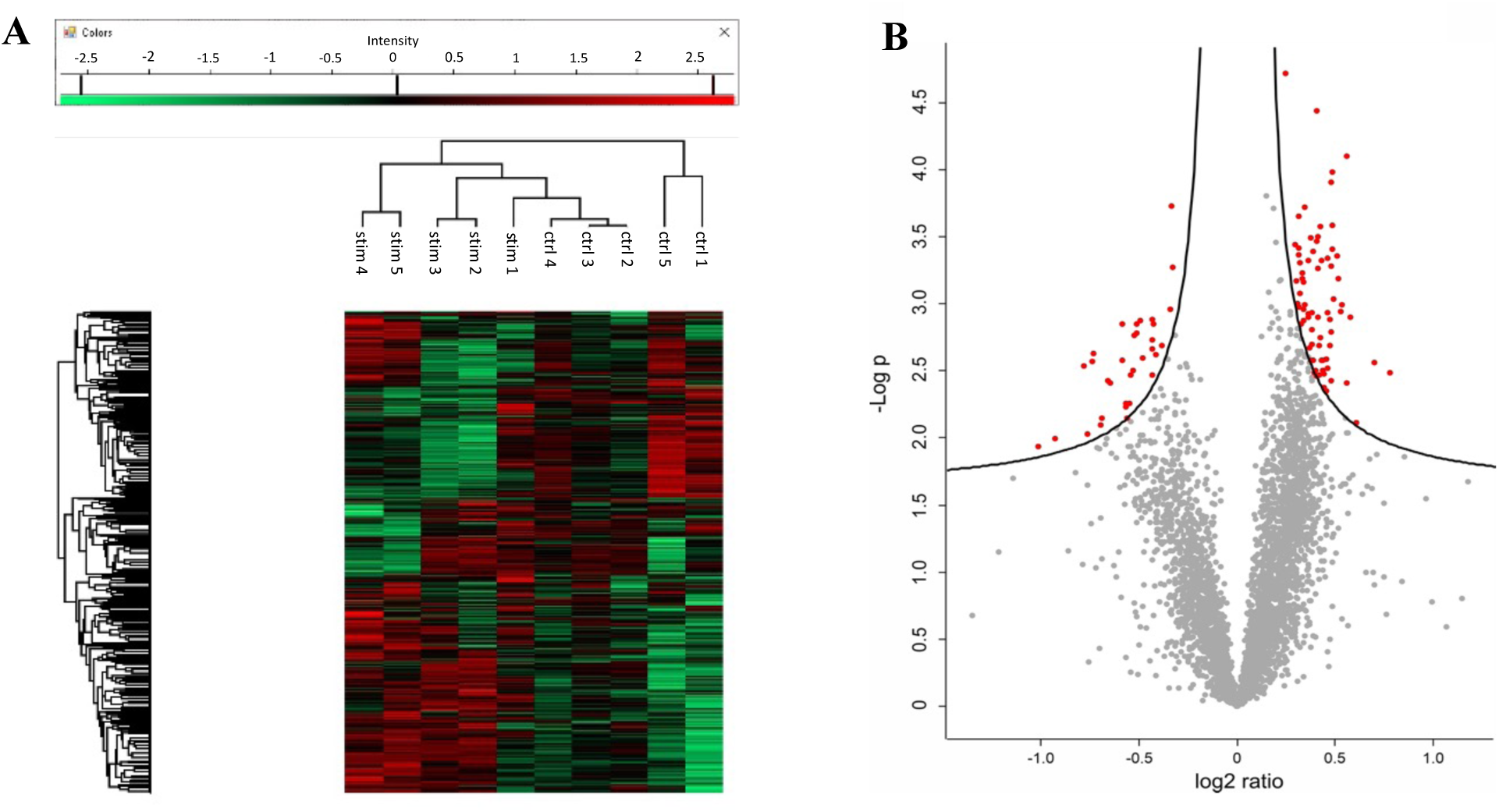
Proteomic assessment of CON-EVs and CCA-EVs. EVs from CON and CCA myotubes were analyzed by LC-MS/MS. (n=5). **(A)** Heatmap showing showing unsupervised clustering of CON-EVs (labelled as ctrl) and CCA-EVs (labelled as stim). Green: decreased expression; red: increased expression. **(B)** Volcano plot depicting significant changes in proteins isolated from CCA-EVs vs. CON-EVs. The red dots indicate proteins that are significantly differential after multiple testing correction. Red dots within the right curve (+log2 ratio) are up-regulated proteins, while those within the left curve (-log2 ratio) are downregulated proteins in CCA-EVs compared to CON-EVs. The gray dots indicate proteins that did not pass the significance threshold (5% false discovery rate).

**Figure 3.**
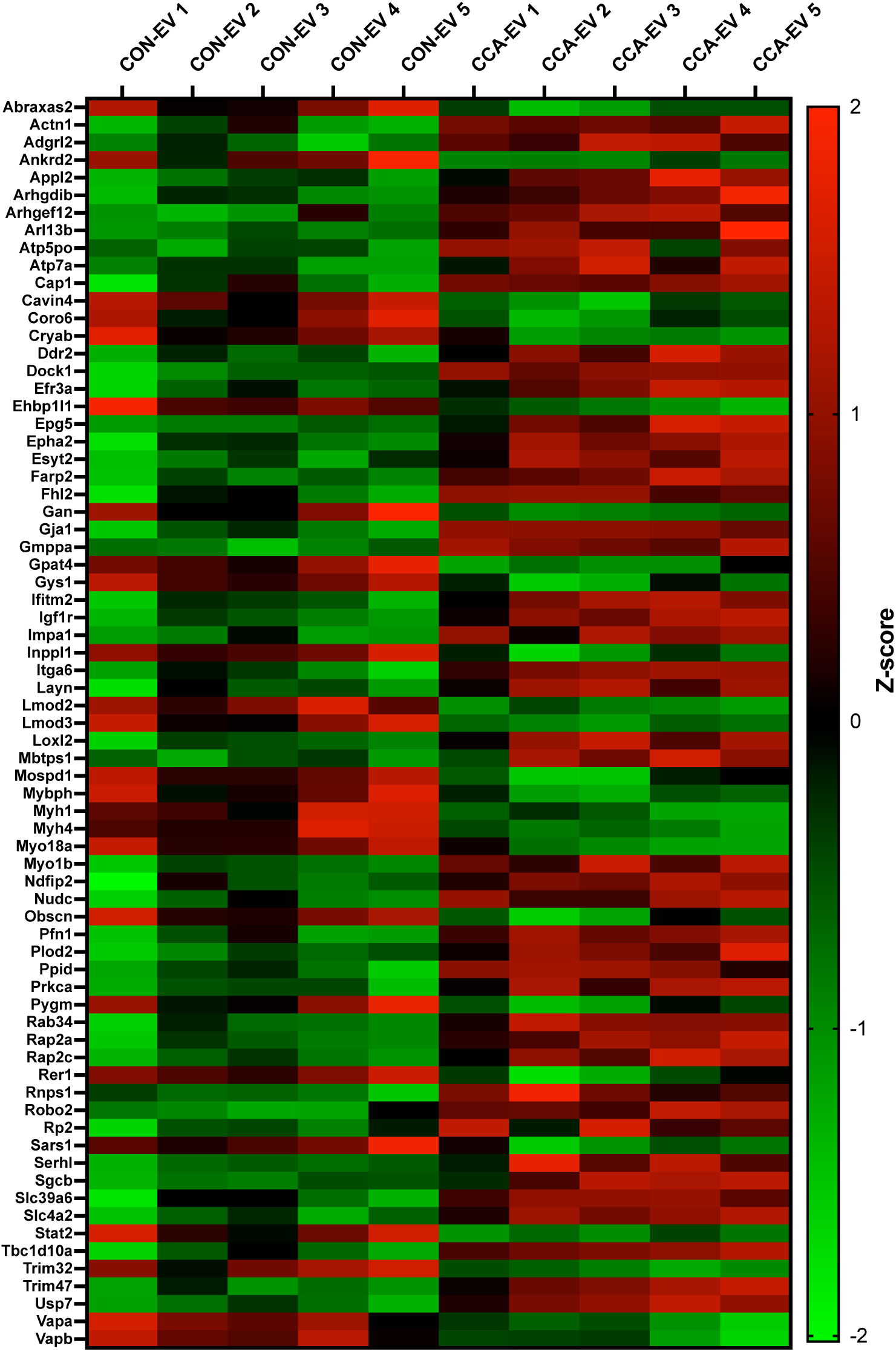
Proteomic analysis of differentially expressed proteins (DEPs). Heatmap of z-scored protein intensities of the DEPs (FDR < 0.05) obtained from five biological replicates comparing CCA-EVs to CON-EVs. Green: decreased expression; red: increased expression. Z-score was calculated as z = (x-µ)/σ using log2 normalized abundance, where x is the raw score, µ is the population mean, and σ is the population standard deviation.

**Table 1.**
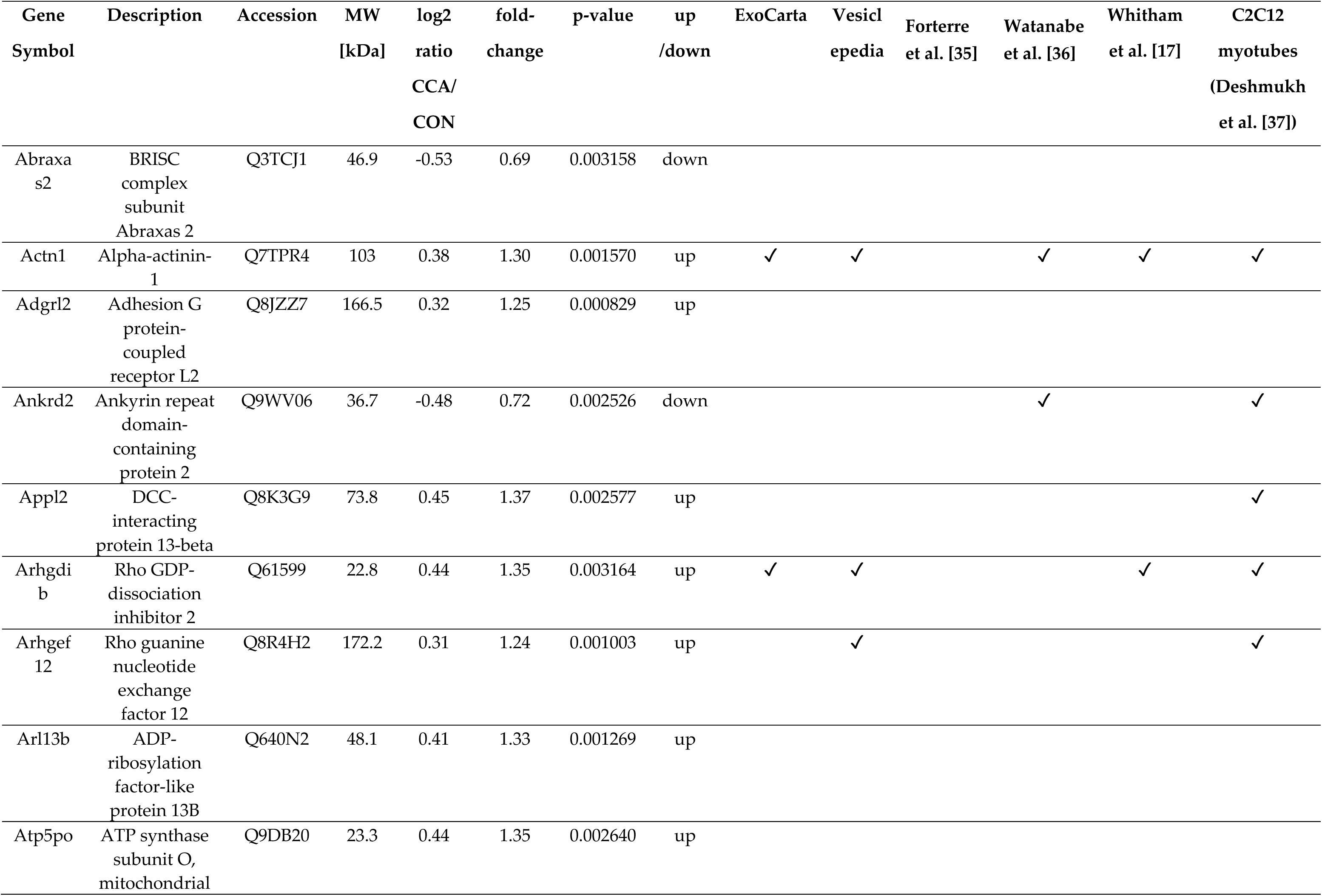
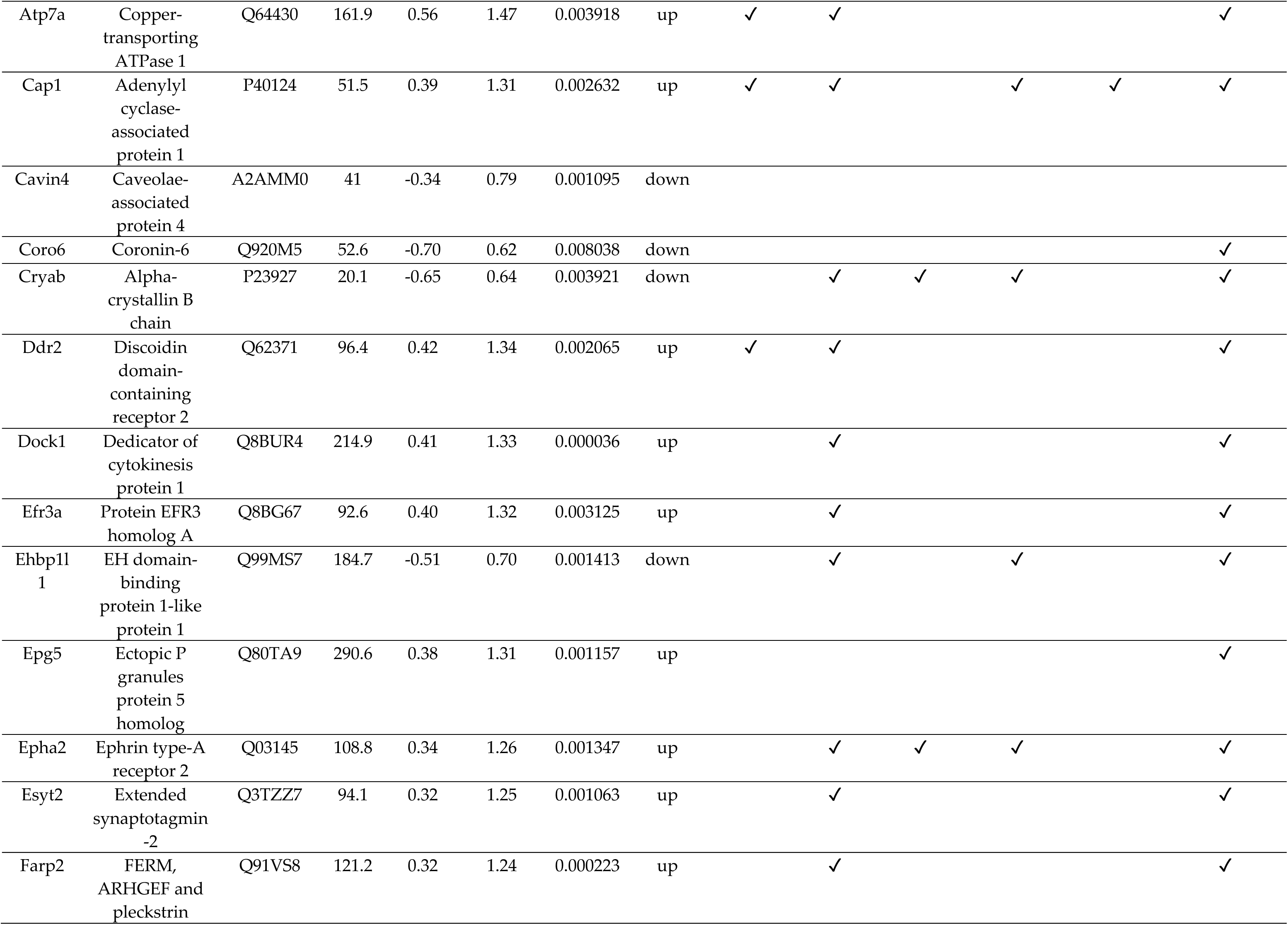
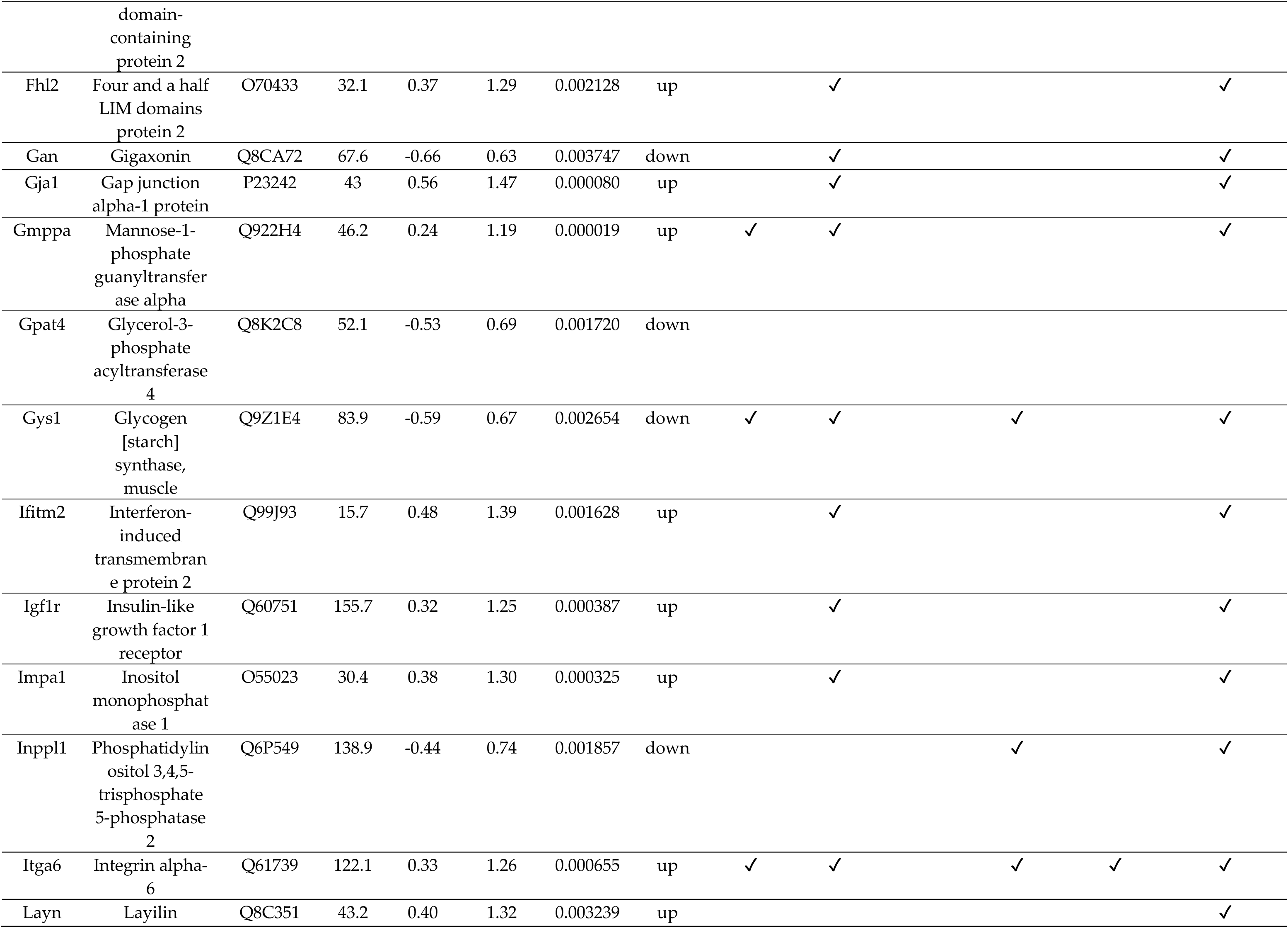
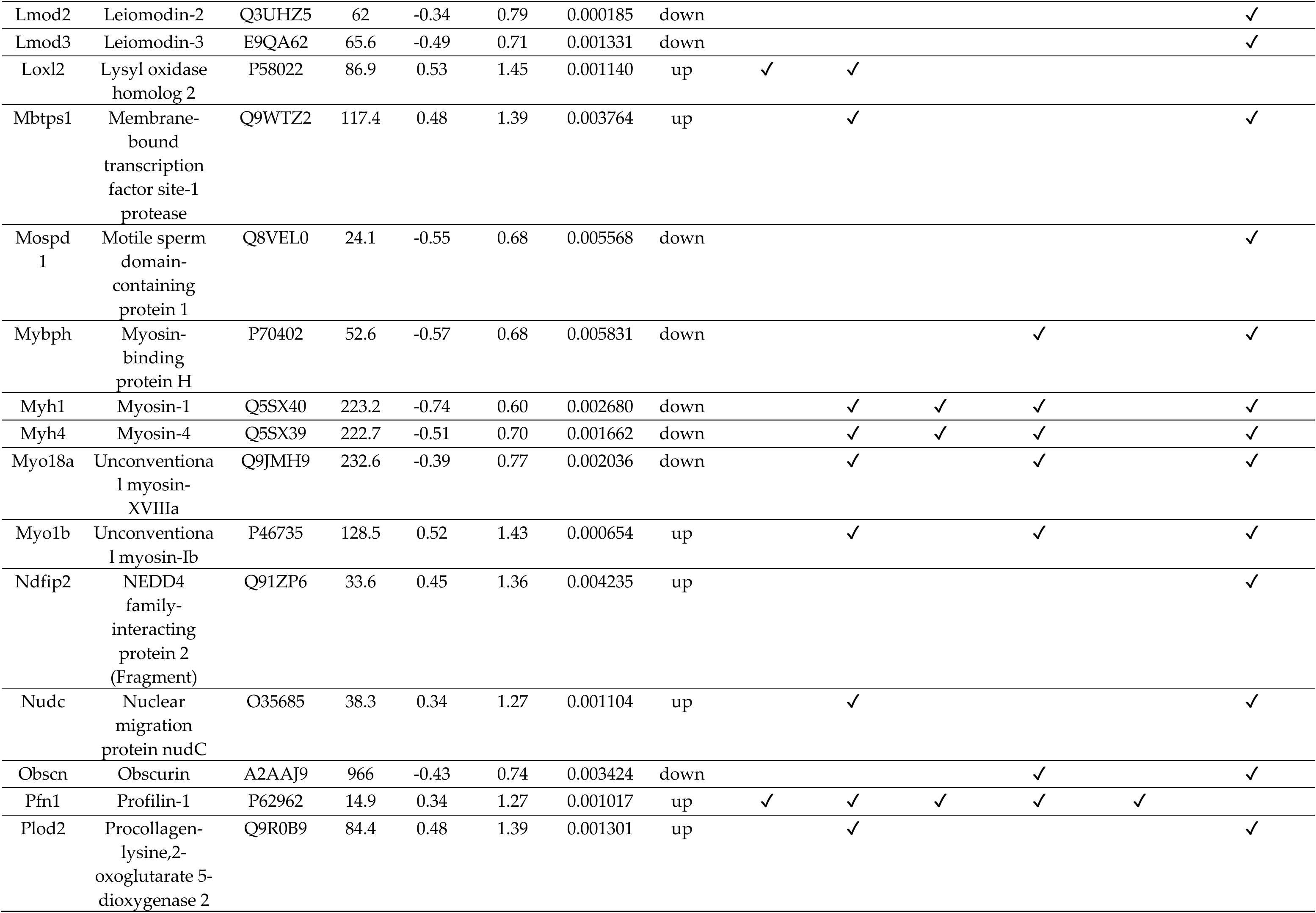
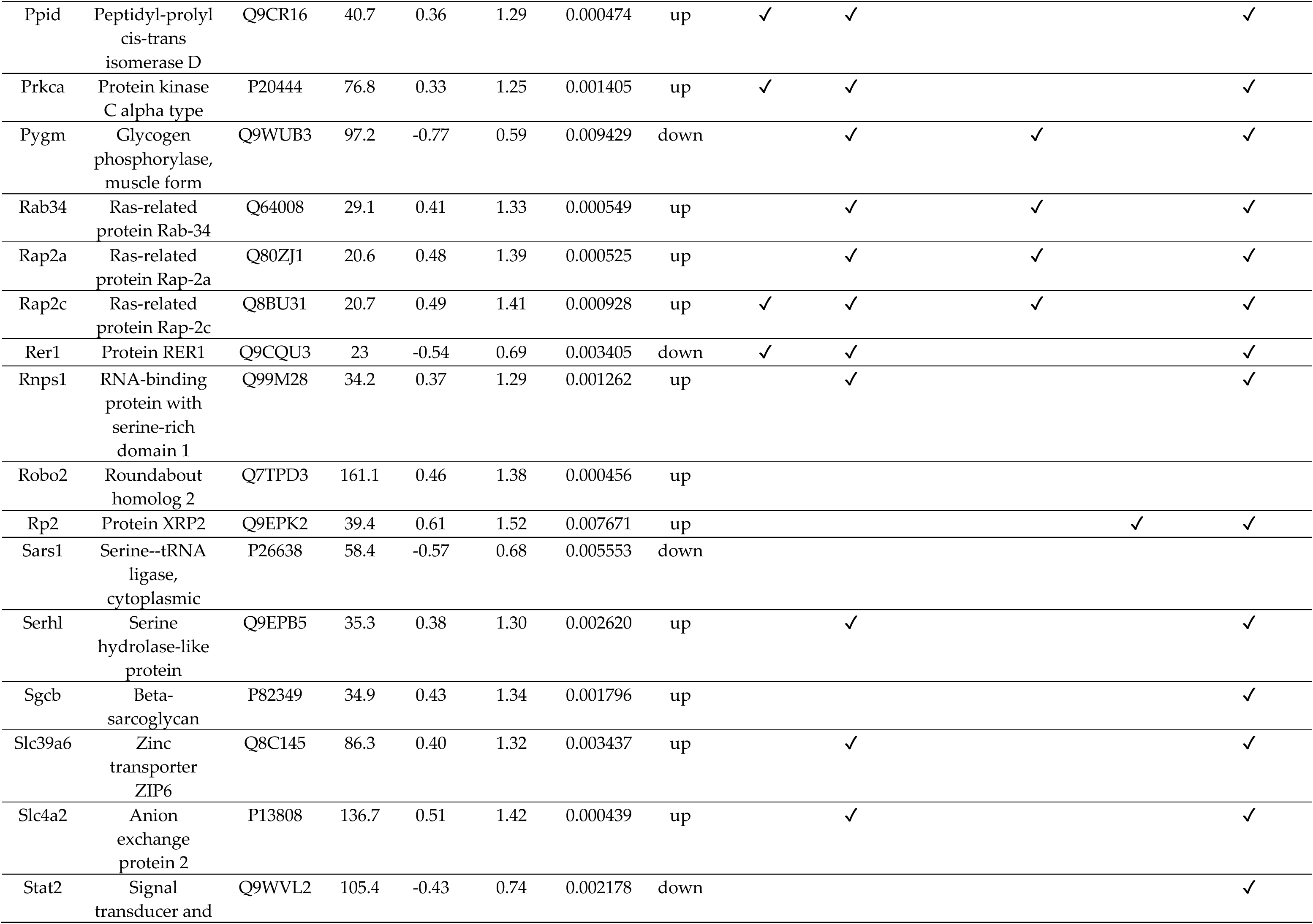
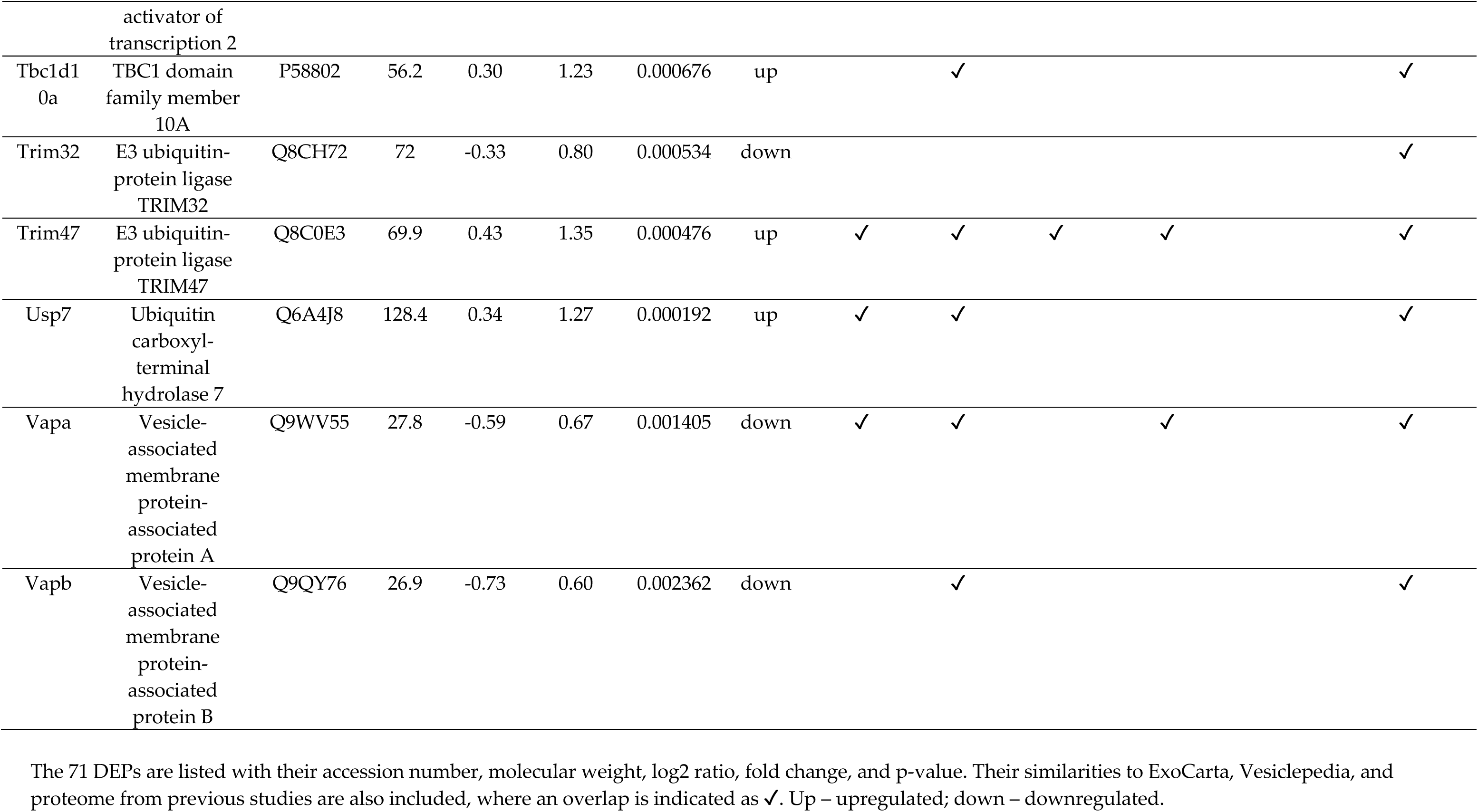
List of differentially expressed proteins in CCA-EVs vs. CON-EVs.

### Comparison of myotube Skm-EV proteome with EV databases

We cross-referenced our dataset with two publicly available EV databases, ExoCarta and Vesiclepedia using FunRich. This analysis showed that 2044 proteins (which is 70% of our 2900 total proteins) have already been previously reported in ExoCarta and Vesiclepedia (**Fig. 4A**). Interestingly, our dataset identifies 856 unique proteins, which have not been listed in any of the EV databases previously (**Fig. 4A** and list of unique proteins in **Table S2**). We also compared the DEPs with the proteins in ExoCarta and Vesiclepedia. Among the 71 DEPs, 24 had not been previously reported, and 47 proteins overlapped with the proteins listed in ExoCarta and Vesiclepedia databases (**Fig. 4B** and **Table 1**). Interestingly, a large proportion of the DEPs that overlapped with the ExoCarta and Vesiclepedia databases were significantly upregulated in CCA-EVs, and belong to protein groups involved in the biogenesis of sEVs. Furthermore, we compared both the total and DEP EV proteome with the top 100 proteins identified on ExoCarta and Vesiclepedia. Among the 2900 proteins, 85 proteins overlapped with the top 100 proteins on ExoCarta, while 84 proteins overlapped with the top 100 proteins on Vesiclepedia (**Fig. 4C**). Importantly, these proteins include sEV markers that are associated with multivesicular body formation (TSG101, PDCD6IP (ALIX), and SDCBP (syntenin-1)), membrane transport and fusion (annexins, RAB proteins and FLOT-1), vesicle adhesion proteins (tetraspanins (CD9, CD81, and CD63) and integrins), and chaperones (heat shock proteins). When comparing the 71 DEPs with the top 100 proteins in both databases, we identified only 4 proteins: PFN1, CAP1, ITGA6, and ACTN1, that overlapped with the top 100 in ExoCarta and Vesiclepedia (**Fig. 4D**).

**Figure 4.**
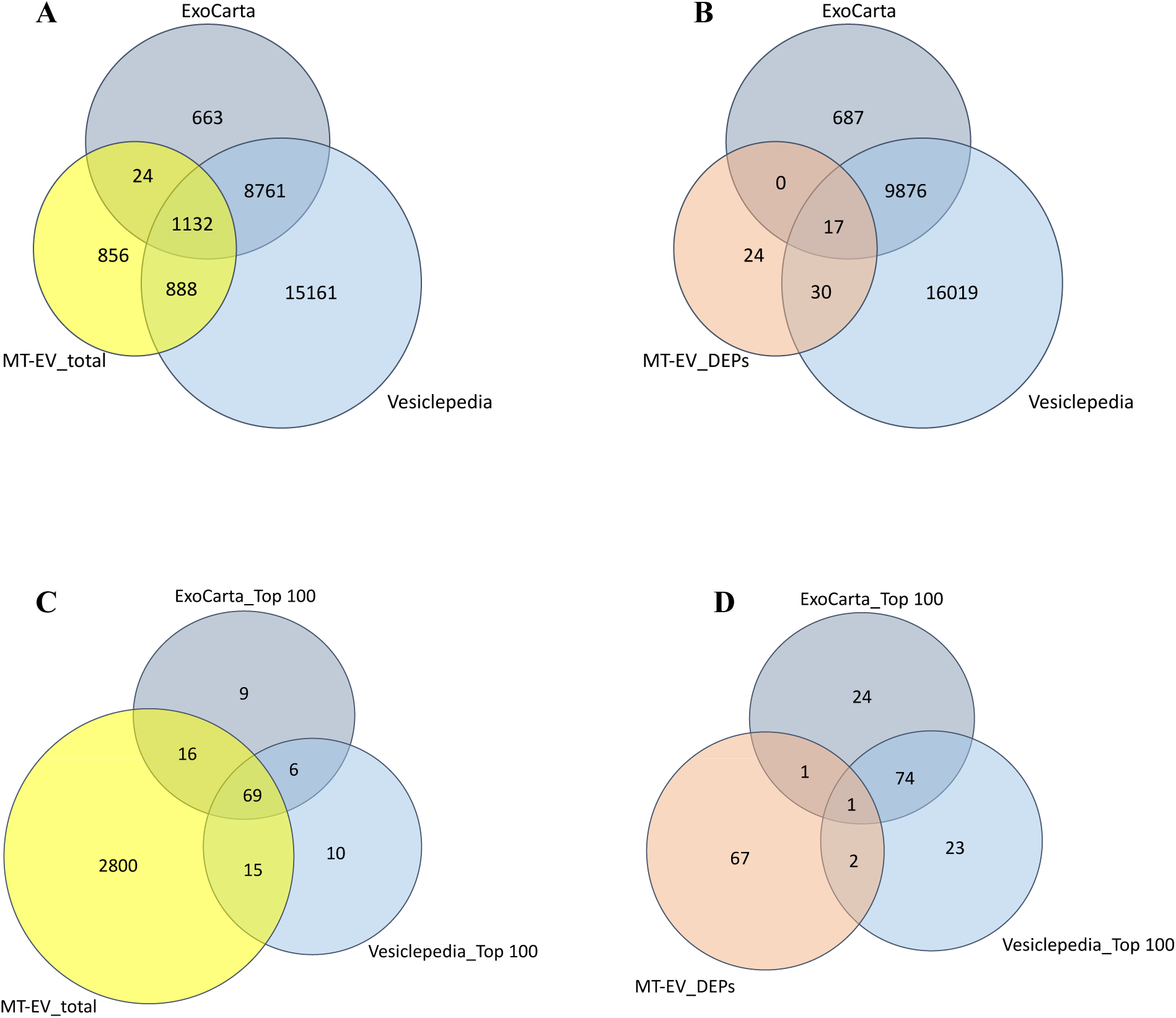
Comparison of EV proteome in present study vs. EV databases. Venn diagrams showing the total proteins (MT-EV_total) and differentially expressed proteins (MT-EV_DEPs) compared with proteins annotated in the ExoCarta and Vesiclepedia databases. Comparison was done between **(A)** MT-EV_total vs. total protein on ExoCarta and Vesiclepedia, **(B)** MT-EV_DEPs vs. total protein on ExoCarta and Vesiclepedia, **(C)** MT-EV_total vs. the top 100 proteins often identified in EVs on ExoCarta and Vesiclepedia, and **(D)** MT-EV_DEPs vs. top 100 proteins identified in EVs on ExoCarta and Vesiclepedia.

### Comparison of myotube Skm-EV proteome with previously published work

Next, we compared the proteins identified in myotube-derived Skm-EVs with and without CCA in our study with two previous studies that have investigated the proteome of myotube-EVs. Forterre et al. and Watanabe et al. identified 383 proteins and 933 proteins in EVs secreted from C2C12 myotubes, respectively [35], [36]. Of the 1316 proteins from the two studies, 838 proteins overlapped with our results, leaving 2062 unique proteins in our study (**Fig. S1A**). When we compared just the DEPs with the 1316 proteins from these two manuscripts, we identified 23 common proteins and 39 unique proteins (**Table 1**). Since there are no previous reports on the proteome of Skm-EVs isolated post-chronic exercise or post-CCA, we compared our dataset with the EV proteome from an exercise study. First we compared our results with Whitham et al., who reported 322 significantly regulated proteins in EVs from an acute bout of exercise vs. rest [17]. Among these, 182 proteins overlapped with our 2900 proteins (**Fig. S1B**), and 6 proteins overlapped with our 71 DEPs (**Table 1**). Thus, we have identified 65 novel DEPs that are putative myokines as they are released from skeletal muscle and modified in response to contractile activity. Lastly, to investigate how specific our EV proteome is to the cell of origin, we compared our dataset with the proteins expressed in the skeletal muscle (using proteome of C2C12 myotubes and mouse triceps muscle from a study by Deshmukh et al [37]). We found that 2656 proteins (92% of the 2900 total proteins in our study) overlapped with proteins expressed in the skeletal muscle, with 244 unique proteins that had not been previously reported (**Fig. 5A**). Since myokines are proteins released by the skeletal muscle in response to exercise, we compared our 71 DEPs with proteins expressed in the skeletal muscle [37]. Remarkably, we found that 61 DEPs overlapped with proteins from the skeletal muscle, with 10 unique DEPs that had not been previously reported (**Fig. 5B** and **Table 1**).

**Figure 5.**
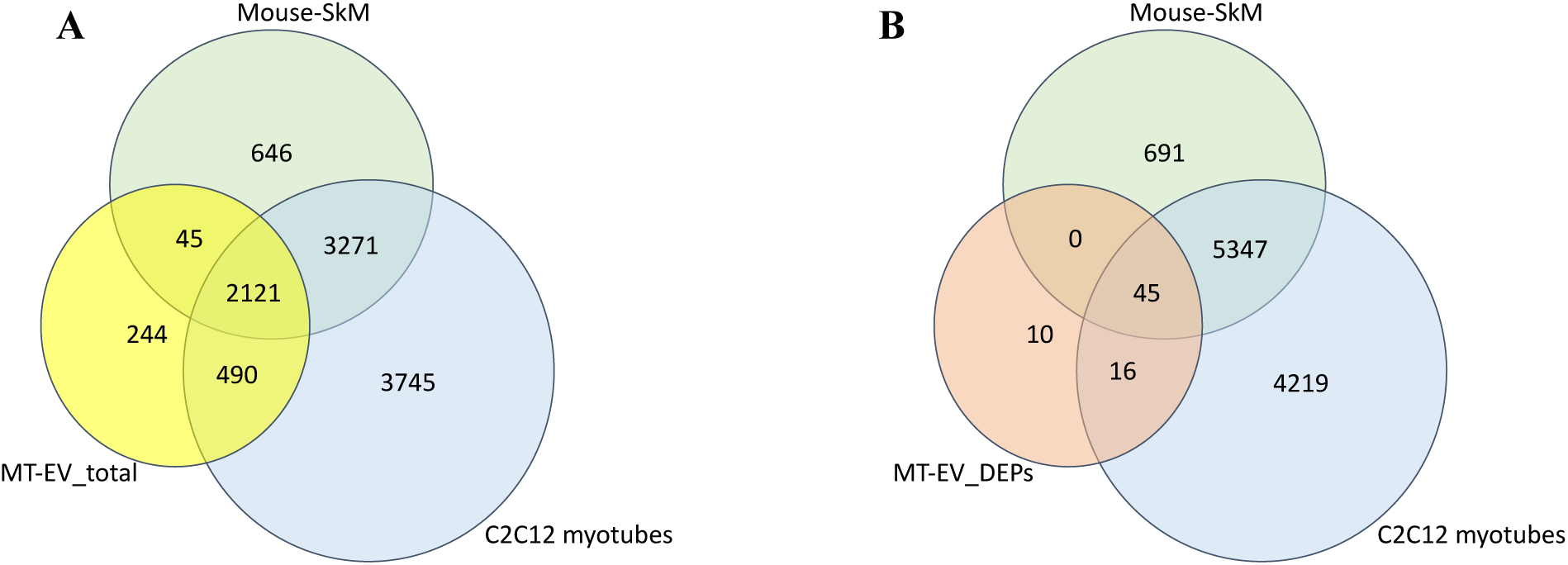
Comparison of EV proteome in present study vs. skeletal muscle proteins. Venn diagrams showing the total proteins (MT-EV_total) and differentially expressed proteins (MT-EV_DEPs) compared with proteins expressed in the skeletal muscle (using proteome of C2C12 myotubes and mouse triceps muscle from Deshmukh et al. 2015 [37]). Comparisons were conducted done between **(A)** MT-EV_total vs. skeletal muscle proteome, and **(B)** MT-EV_DEPs vs. skeletal muscle proteome.

### Functional enrichment analysis of DEPs

To obtain further insight into the subcellular origin and function of the DEPs, we did a functional enrichment analysis using FunRich. The analysis showed that 55.6% (p<0.001) of the DEPs originated from the cytoplasm, and 45.4% from the plasma membrane (p=0.001, **Fig. 6A**). The top molecular function of the DEPs is actin filament binding (11%, p<0.001; **Fig. 6B**). The top biological processes that the DEPs were involved in included actin filament organization (12.1%, p<0.001), muscle contraction (5.6%, p=0.018), and positive regulation of protein secretion (5.6%, p=0.054, **Fig. 6C**).

**Figure 6.**
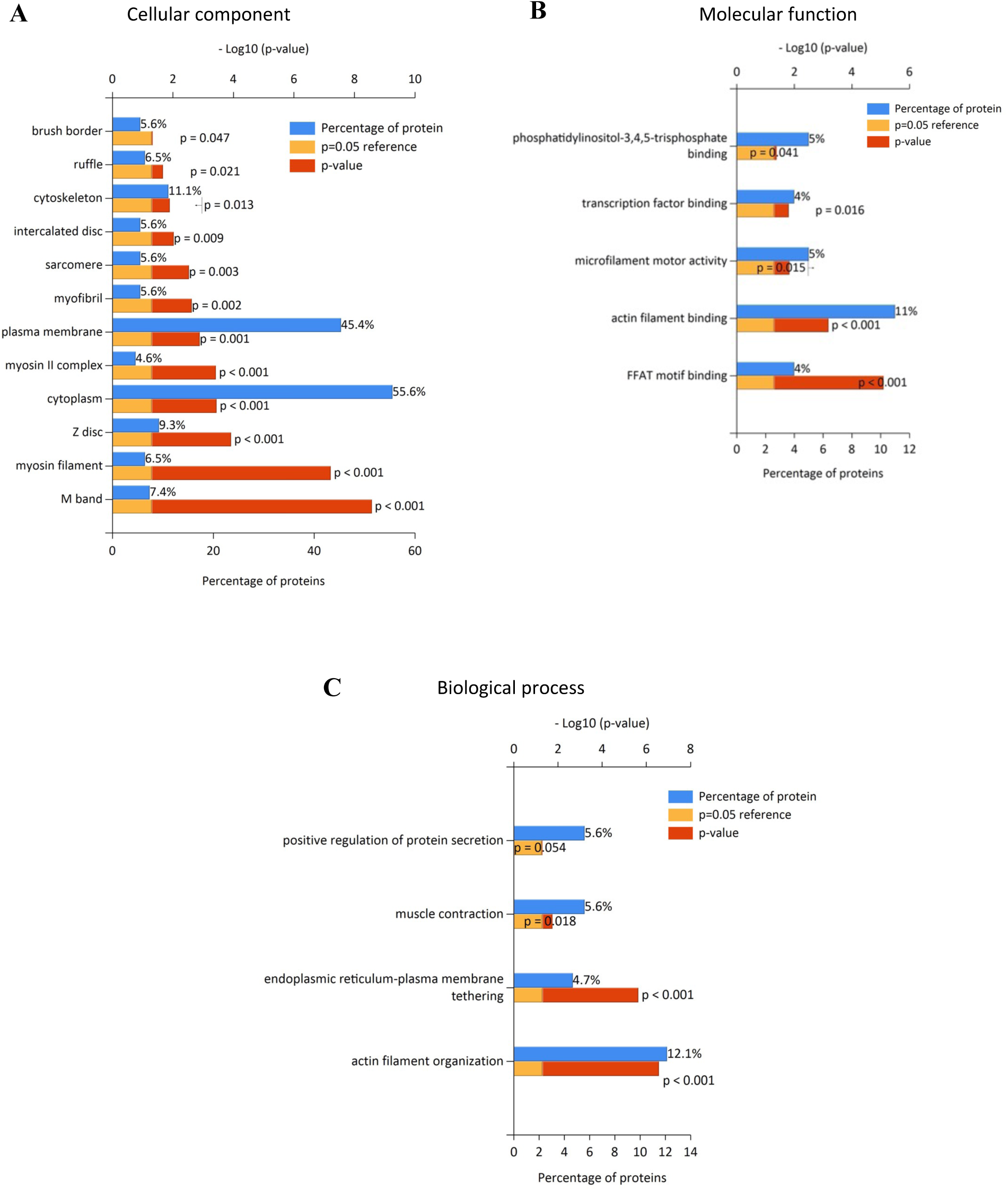
Functional enrichment analysis of differentially expressed proteins. Functional enrichment analysis of the DEPs was performed using the FunRich software for **(A)** cellular component, **(B)** molecular function, and **(C)** biological process when results were ≥ 4% or higher for protein percentage associated with each. Blue bars represent the percentage of proteins assigned to the indicated term, yellow bars show the reference p value (0.05), and red bars show the calculated p value of enrichment for the indicated term.

### Pathway analysis of DEPs

To perform a network analysis, we uploaded the DEPs to the STRING database to identify interactions between them. In total, there were 71 nodes (DEPs) and 43 edges (predicted functional associations) observed in the network. The most enriched network cluster in STRING was associated with muscle protein and myofibril assembly, and striated muscle contraction and sarcomere organization (**Fig. 7A**). A KEGG analysis identified the top three enriched pathways based on -log10(FDR) values including focal adhesion, Rap1 signaling pathway and regulation of actin cytoskeleton (**Fig. S2**). To further identify signaling networks, we conducted an IPA analysis with the 71 DEPs. IPA identified some canonical pathways, of which the top two pathways included actin cytoskeleton signaling and integrin signaling (**Fig. 7B**). IPA also identified significant putative biological functions that could be altered by DEPs such as cellular movement, cell death and survival (data not shown). In particular, IPA analysis revealed networks associated with several mediators such as CaMKII, pdgf complex, PDGF BB, Akt, Phosphoinositide 3-kinase (PI3K), cAMP Response Element-Binding Protein (CREB), Osteocalcin, Integrin, c-Src and Filamin (**Fig. 8**).

**Figure 7.**
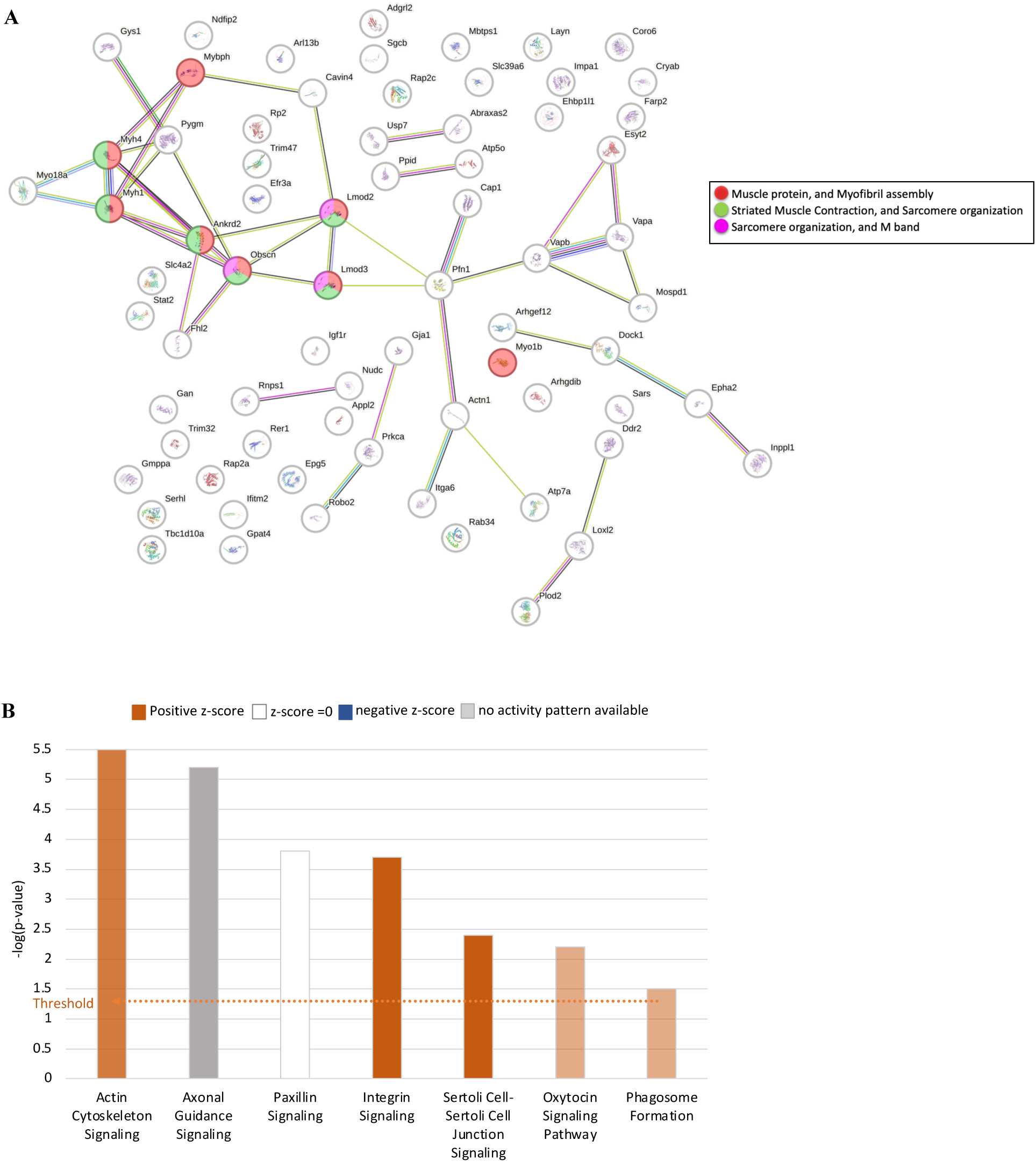
Pathway analysis of differentially expressed proteins. **(A)** Interaction networks of DEPs was built using the STRING database. Colors of nodes designate involvement in the assigned local network cluster: Red – muscle protein and myofibril assembly; green – striated muscle contraction and sarcomere organization; and pink – sarcomere organization and M band. **(B)** The top canonical pathways affected with the DEPs are identified by IPA with −log10 (p-value) and z-score indicated. Orange bars: positive z-score; blue bars: negative z-score; grey bars: no activity pattern available; white bars: z-score = 0. The darker coloring of the bars represents the more significant z-scores.

**Figure 8.**
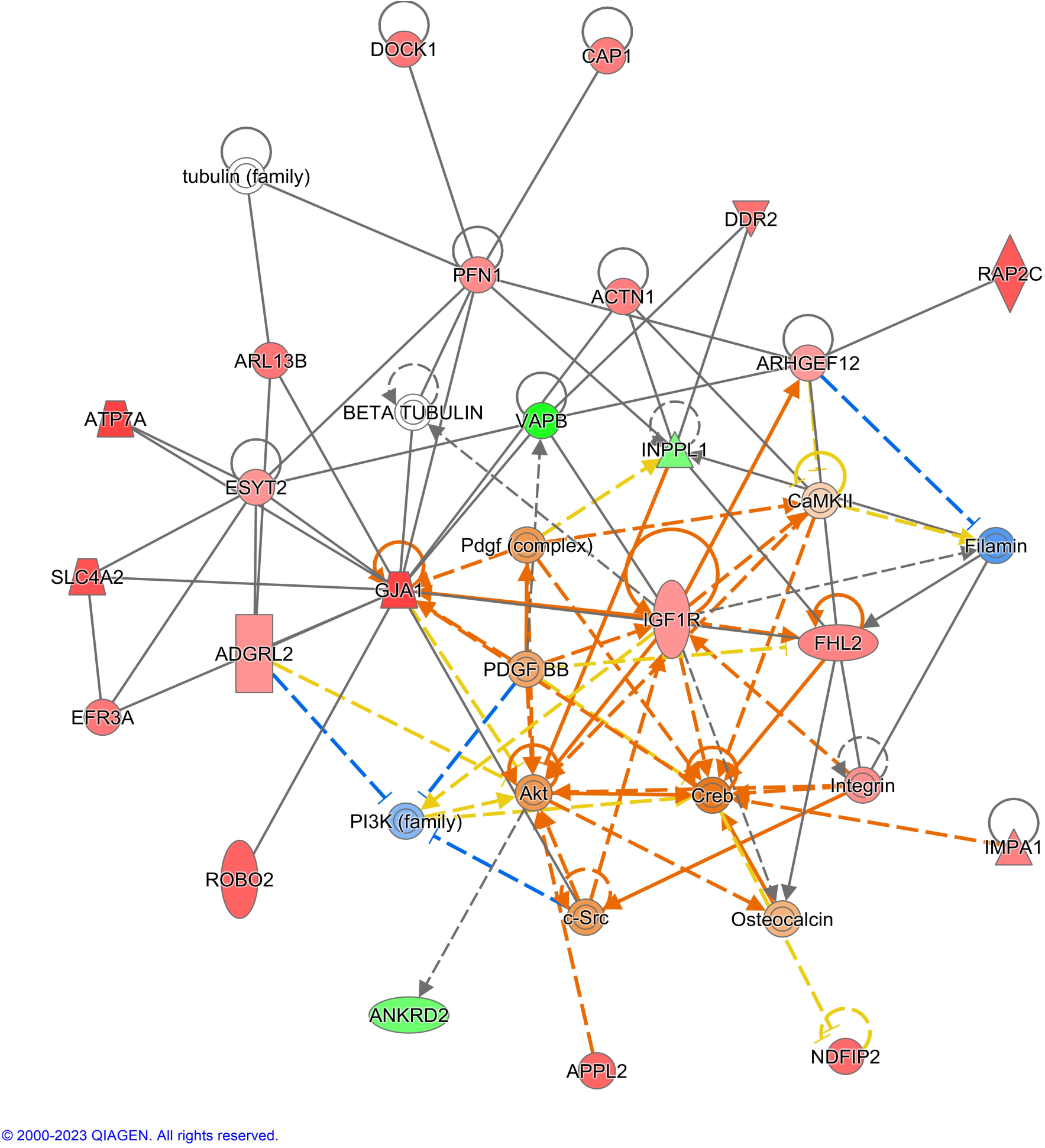
Functional network analysis associated with differentially expressed proteins. Network analysis using the IPA software was conducted using selected DEPs to predict downstream effects, including identification of putative targets and building interactive signalling networks. Solid line - direct interaction; Dotted line - indirect interaction. Red – upregulated DEPs; green – downregulated DEPs (according to color intensity). Other colours represent proteins not contained in the present data: orange - predicted activation; blue – predicted inhibition; yellow - inconsistent findings; and gray - no prediction can be made.

### Relevance scoring and validation of DEPs by western blotting

The DEPs were given a relevance score based on their function (i.e., associated with mitochondrial function or inhibition of cancer growth/metastasis), subcellular location (membrane-bound), and fold change (>1.25). Importantly, TMT-based quantitation is known to suffer from ratio compression such that actual fold-changes are typically higher than determined by TMT [38]. Using these criteria, we generated a relevance score table, and proteins that met the criteria were assigned a score of “2” for each criterion, while those that did not were given a score of “1”. The total scores were then calculated, with nine proteins scoring ≥7. From these, the top five were selected for initial investigation, with remaining to be examined in future studies. The five DEPs with the highest scores - GJA1 (Gap junction alpha-1 protein) also known as Connexin 43, PFN1 (Profilin-1), PRKCA (Protein kinase C alpha type), IGF1R (Insulin-like growth factor 1 receptor), and ATP7A (Copper-transporting ATPase 1) - were chosen for validation. The complete list of DEPs with their relevance scores can be found in **Table S3.** Additionally, we selected adhesion protein, ITGA6 (Integrin alpha-6), for validation based on data from Whitham et al. [17] as discussed below. All six proteins were validated using western blot with four independent biological samples. Interestingly, the expression level of all the six proteins were upregulated in CCA-EVs by ∼2-fold compared to CON-EVs (**Fig. 9A** and **9B**), confirming the results obtained from the proteomic analysis (**Fig. 9C**). This is in agreement with the abovementioned ratio compression of TMT datasets. To investigate the location of these six proteins in EVs, we treated CCA-EVs with Triton with or without Pro K before immunoblotting. The expression of ITGA6, IGF1R, ATP7A, and PFN1 was abolished when EVs were pre-treated with Pro K (**Fig. 9A**). Only GJA1 and PRKCA remained intact, albeit in reduced levels, with Pro K treatment (**Fig. 9A**). The expression of all six proteins remained intact when EVs were permeabilized with Triton; however, GJA1, IGF1R, and PFN1 levels were reduced. Finally, none of the six proteins were detected when CCA-EVs were co-treated with Triton and Pro K (**Fig. 9A**).

**Figure 9.**
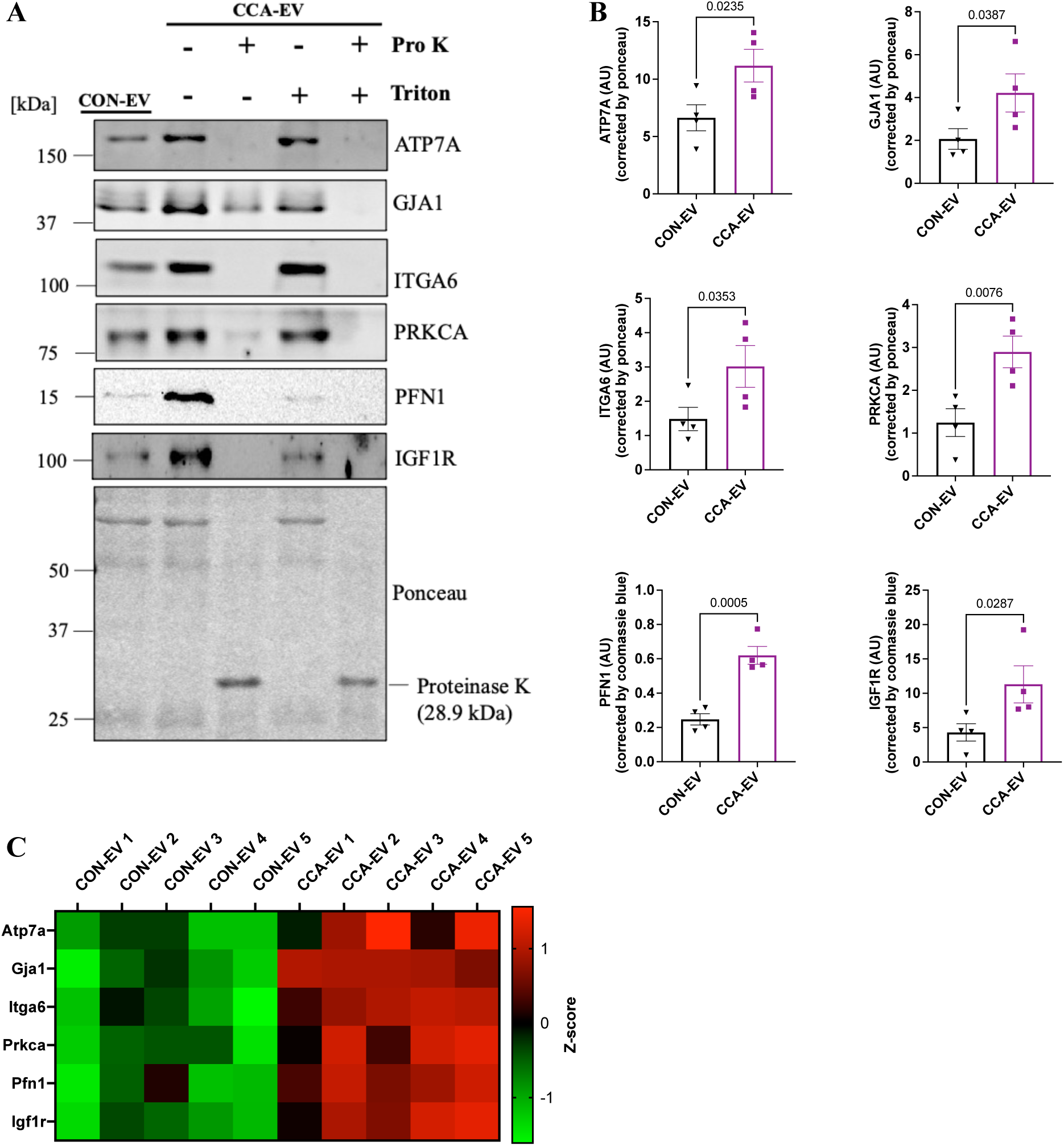
Validation of DEP targets identified through proteomics in CON-EVs and CCA-EVs. Isolated EVs after each bout of CCA (days 1-4) were pooled together, proteins isolated and subjected to immunoblotting. CCA-EVs were also pretreated with Triton X-100 with or without proteinase K before protein extraction and Western blotting. **(A, B)** Representative western blots and quantification of ATP7A (∼180 kDa), GJA1 (39-44 kDa), ITGA6 (125-150 kDa), PRKCA (80 kDa), PFN1 (15 kDa) and IGF1R (95 kDa). **(C)** Heatmap of z-scored protein intensities showing expression of selected DEPs in CCA-EVs compared to CON-EVs. Data were analyzed using an unpaired Student’s t-test and expressed as scatter plots with mean ± SEM error bars (n=4). Exact p values for significant results (p<0.05) are shown.

## Discussion

To evaluate the proteome of Skm-derived EVs, we performed a comprehensive LC-MS/MS analysis of EVs isolated from control and chronically stimulated C2C12 myotubes. For the CCA model, we used our established protocol optimized to mimic chronic endurance exercise *in vitro* [27]. We have previously shown that CCA increased the concentration of myotube-EVs, and these in turn increased mitochondrial biogenesis in C2C12 myoblasts [27], and reduced cell viability, induced apoptosis, and senescence in non-small cell lung cancer cells [28]. However, the underlying mechanisms remain unknown. In this study, we investigated the proteome of CON-EVs and CCA-EVs and identified several putative proteins and signaling pathways that may be critical in mediating the beneficial effects perpetuated by CCA-EVs.

Using a LC-MS/MS-based proteomics profiling approach, we identified a total of 2900 proteins in Skm-EVs from murine myotubes, where 71 proteins were differentially expressed between CON-EVs and CCA-EVs. Of the 71 proteins, 46 were significantly upregulated while 25 were significantly downregulated in CCA-EVs. We compared our findings to the EV databases, ExoCarta and Vesiclepedia, which include all proteins identified in EVs from different types of tissues or cells, isolation methods, and detection methods. We were able to determine an overlap of 2044 proteins between our results and previously documented EV proteins in ExoCarta and Vesiclepedia, illustrating a common protein profile of EV regardless of the type of cell, method of isolation or characterization. Among the DEPs, we identified an overlap of 47 proteins with proteins listed on the two EV databases, of which a large proportion were upregulated in CCA-EVs. Furthermore, we identified 4 proteins that overlapped with the top 100 proteins on ExoCarta and Vesiclepedia including PFN1, CAP1, ITGA6, and ACTN1. Interestingly, these proteins form a network (predicted functional associations) and are upregulated in CCA-EVs. Additionally, we compared our findings to previous reports of changes in EV proteome with exercise. Whitham et al. showed that acute exercise significantly altered the expression of 322 proteins, notably including PFN1, CAP1, ITGA6, and ACTN1 [17]. Altogether our data is in line with previous reports, and identifies new unique and novel proteins associated with CCA-EVs. The four proteins common to both our work and Whitham et al. [17], should be noted as findings of importance that warrant further research. The DEPs could be playing an important role in the increased secretion of CCA-EVs from murine myotubes [27], and/or involved in inducing downstream functional effects [27], [28].

Next, we compared our data to previous studies that have investigated the proteome of myotube-EVs [35], [36], our current study revealed 2062 proteins that have not been identified in other myotube-EVs studies before. The discovery of these unique EV proteins could be because studies investigating Skm-EVs are sparse. To the best of our knowledge, no one has investigated the effect of chronic exercise on the proteome of Skm-EVs specifically. Studies that have investigated EV proteome following acute exercise have isolated EVs from plasma [17], [39], which would comprise a heterogenous mixture of vesicles derived from platelets, erythrocytes, endothelial cells, leukocytes, [18], [40], and with a comparatively little proportion of EVs originating from skeletal muscle [41], [42]. Since EVs are a snapshot of the cells that released them, we investigated how specific our dataset is to the proteome of skeletal muscle. We compared our dataset to proteins expressed in C2C12 myotubes as well as mouse skeletal muscle (triceps muscle) [37]. Interestingly, 92% of our total protein overlapped with the proteins from C2C12 myotubes and mouse skeletal muscle, which demonstrates the importance of this current work in evaluating the Skm-EVs specifically. Given the challenges in distinguishing Skm-EVs from systemic EV preparations, and the fact that skeletal muscle is the primary organ involved in muscle contraction and the release of myokines during physical activity, it is critical to distill the impact of chronic exercise on the characteristics of Skm-EVs specifically. The CCA model allows us to circumvent the logistical difficulties in purifying skeletal muscle EVs from circulation. The large similarity in the CCA-EV proteome in our study to previous reports of mouse skeletal muscle proteins highlights the relevance and validity of using this experimental approach to interrogate skeletal muscle-derived EVs.

Myokines are proteins released by the skeletal muscle in response to exercise, thus we sought to identify potential myokines in our data. First, we searched the literature to identify bona fide myokines, which we manually cross-referenced with our dataset. The DEPs in our study did not contain any of the published myokines. However, among the non-significantly regulated proteins in our data, we found canonical myokines such as decorin, secreted protein acidic and rich in cysteine (SPARC), follistatin, follistatin related protein (FSTL1), and some associated proteins including oncostatin-M-specific receptor subunit beta (OSMR), leukemia inhibitory factor receptor protein (LIFR) and interleukin-6-receptor subunit beta (IL6ST) [9], [11], [43]. To identify putative novel myokines in our data, we compared the 71 DEPs with proteins expressed in the skeletal muscle, and found an overlap of 61 DEPs. Since, these 61 proteins are expressed in the skeletal muscle, significantly altered with chronic contractile activity and secreted encapsulated within EVs, we classified them as putative myokines. Using a similar approach, Whitham et al. previously identified 35 novel candidate myokines in plasma-EVs isolated from the femoral vein of humans following an acute bout of exercise [17]. Interestingly, none of the candidate myokines from Whitham et al. were present in our dataset, which indicates that we identified 61 novel potential myokines. Further research into the endocrine, autocrine and/or paracrine functions of these novel putative myokine candidates is needed. Overall, our findings support EVs as an alternate pathway for myokine secretion.

Functional enrichment analysis showed that the top cellular components of the EV DEPs were cytoplasm and plasma membrane, which are common sites for biogenesis of exosomes as well as microvesicles [14]. Unsurprisingly, the functional analysis results using FunRich, KEGG, STRING and IPA revealed that the top enriched pathways involve actin filament binding and organization, integrin signaling, and muscle contraction, which are important pathways required for the proper functioning of the skeletal muscle [44], [45]. Interestingly, actin polymerization is one of the processes known to be involved in the biogenesis of EVs [46]. Furthermore, studies have shown that adhesion proteins such as integrins on the surface of the vesicles are likely responsible for the target specificity of EVs [46]. In this current study, we observed a significant increase in a wide range of proteins involved in cell adhesion in CCA-EVs, including ITGA6, ACTN1, DOCK1, PFN1, and RAP2A. In concurrence with our findings, Whitham et al. reported a significant release of adhesion protein, ITGB5, after acute exercise, alongside an increased uptake of this protein in liver cells treated with EVs from exercised mice [17]. Additionally, Fuentes et al. reported that ITGB3-mediated uptake of sEVs facilitates intercellular communication in breast cancer cells [47]. The results from these two studies prompted us to ask the question “Could ITGA6 or any of the adhesion proteins be involved in the specificity and uptake of CCA-EVs into recipient cells?” Therefore, we selected ITGA6 as one of the proteins to be validated by western blotting, in addition to the top five selected by the relevance scoring process, as discussed below.

Apart from ITGA6, we selected five other candidate DEPs for validation based on relevance scoring results. We had recently demonstrated that CCA-EVs increased mitochondrial content, function, and respiration in C2C12 myoblasts [27], and decreased cell viability, induced apoptosis, and evoked senescence in non-small cell lung cancer cells [28]. The functional effects of CCA-EVs were abolished when CCA-EVs were pre-treated with Triton with or without Pro K [27], [28]. Pre-treatment of EVs with Pro K digests all exposed proteins on EV membrane, and in the EV corona, leaving the proteins in EV lumen intact. With Triton, EV membrane would be permeabilized, and in the combination group of both Triton followed by Pro K, the digestion of all EV proteins both outside and inside the EV membrane will be affected. Therefore, our results indicated that the adaptive effects induced by CCA-EVs in recipient cells were likely mediated by the membrane-bound EV protein cargo [27], [28]. With this in mind, we selected candidate upregulated DEPs based on their function (i.e., associated with mitochondria function and/or inhibition of cancer growth), subcellular location (membrane-bound), and fold change (>1.25). From the relevance score, the five of the top nine highest-scoring DEPs were selected for initial investigation including GJA1, PFN1, PRKCA, IGF1R, and ATP7A. Interestingly, all five proteins have been previously identified as EV-associated proteins on Vesiclepedia. All five proteins are also involved in regulating mitochondrial function, turnover, biogenesis and/or dynamics. GJA1, also known as connexin 43, localizes to the inner and outer mitochondrial membranes, binds to microtubules to promote mitochondrial transport and enhance mitochondrial movement, induces mitochondrial fission, and protects mitochondria from oxidative stress [48]. Like GJA1, PFN1 also localizes inside mitochondria specifically within the matrix, and regulates mitochondrial morphology, dynamics, and respiration [49]. The activation and mitochondrial translocation of PRKCA (or PKCα) is necessary for improving mitochondrial function after ischemic injury. This occurs by regulating the levels of critical subunits in the electron transport chain complexes and ATP synthase, as well as modulating mitochondrial levels of sirtuin-3 (SIRT3) and sirtuin-5 (SIRT5) [50]. IGF-1 signaling has been shown to increase transcription of proteins involved in mitochondrial biogenesis such as PPAR gamma co-activator-1 beta (PGC-1ß), and also regulates mitochondrial dynamics and turnover through the activity of Nuclear factor erythroid 2-related factor 2 (NRF2) and BCL2/adenovirus E1B 19 kDa protein-interacting protein 3 (BNIP3) [51]. ATP7A transport activity is essential in maintaining the redox balance in the mitochondria by protecting it from excessive copper entry [52]. Furthermore, from IPA analysis, we observed that some of these selected proteins are associated with several signaling proteins including Creb and Akt, which are linked to mitochondrial biogenesis [53], [54]. GJA1 and PFN1 have also been reported to be involved in the inhibition of cancer growth [55], [56]. It is possible that one or more of these five DEPs are involved in the pro-metabolic, and anti-tumorigenic effects induced by CCA-EVs, we reported recently [27], [28].

After the selection of the candidate DEPs, we validated their expression in CON-EVs and CCA-EVs using western blotting. Notably, the western blot analysis, showed that all selected six proteins (including ITGA6) were upregulated by ∼ 2-fold in CCA-EVs compared to CON-EVs, confirming the results obtained from the proteomic analysis. To investigate the location of these six proteins in EVs, we treated CCA-EVs with Triton with or without digestion with Pro K before immunoblotting. The expression of ITGA6, IGF1R, ATP7A, and PFN1 was abrogated with Pro K treatment, indicating that these proteins were either membrane-associated and/or in the EV corona. The expression of GJA1 and PRKCA was reduced with Pro K treatment, indicating the presence of these proteins in the lumen of EVs. We also observed that all six candidate DEPs remained detectable post-Triton treatment, albeit three exhibited reduced abundance. However, it is important to note that presence of these EV proteins after Triton treatment does not indicate functional activity in recipient cells. In addition to perpetuating functional effects, the DEPs could also be involved in EV uptake. An important mechanism of EV uptake is the binding of EVs via specific surface receptors to the recipient cell, which co-incidentally triggers various intracellular signaling pathways that modulate recipient cell function. The binding of EVs to target cells is determined by several factors including the composition of the EV membrane, such as transmembrane/membrane-bound proteins and lipids [57]. This leads us to speculate that the proteins on the surface of CCA-EVs (e.g., adhesion proteins such as ITGA6 and PFN1) may be required for EV targeting and internalization through multiple pathways, and the removal of these proteins likely blocks the delivery of EV cargo into recipient cells. Altogether, the data suggest that the DEPs in CCA-EVs likely contribute to mediating the downstream beneficial effects of EVs and/or could also be involved in EV-receptor binding in recipient cells. Further studies are required to confirm whether enhanced cargo delivery in recipient cells is in line with the DEPs identified, and delineate the specificity of the target proteins and downstream signaling pathways that are activated by the DEPs in the CCA-EV cargo.

In summary, our data provide a unique and novel insight into the proteome of Skm-EVs derived post-CCA. We have identified novel putative myokines and potential protein candidates that may mediate the beneficial effects of CCA as transmitted by EVs which we recently reported. Functional assays (e.g., knockdown or overexpression of proteins) to delineate which specific protein(s) are involved in perpetuating the pro-metabolic and anti-cancerous effects of CCA-EVs are needed. The results may lead to the identification of targets that can be therapeutically leveraged in conditions characterized by poor metabolic capacity, and in cancer therapeutics.

## Author Contributions

P.O.O., Y.L., and T.F.G.S. performed experiments in the current study, analyzed data, and created figures. P.O.O. and A.S. helped write and revise the manuscript. K.J.M., J.W.G., R.L, and R.P.Z. provided technical and theoretical expertise to complete the work. All authors were involved in manuscript revisions. A.S. designed the project, and helped synthesize data, create figures, write, and edit the manuscript. A.S. is the corresponding author and directly supervised the project. All authors have read, edited and agreed to the published version of the manuscript.

## References

[1] R. G. C.S. Adeel, and A. Zolt, “Running Forward,” Circulation, vol. 129, no. 7, pp. 798–810, Feb. 2014, doi: 10.1161/CIRCULATIONAHA.113.001590.

[2] J. Viña, F. Sanchis-Gomar, V. Martinez-Bello, and M. C. Gomez-Cabrera, “Exercise acts as a drug; The pharmacological benefits of exercise,” British Journal of Pharmacology, vol. 167, no. 1. Wiley-Blackwell, pp. 1–12, Sep. 2012, doi: 10.1111/j.1476-5381.2012.01970.x.

[3] G. N. Ruegsegger and F. W. Booth, “Health Benefits of Exercise,” Cold Spring Harb. Perspect. Med., vol. 8, no. 7, Jul. 2018, doi: 10.1101/CSHPERSPECT.A029694.

[4] M. A. Febbraio, N. Hiscock, M. Sacchetti, C. P. Fischer, and B. K. Pedersen, “Interleukin-6 is a novel factor mediating glucose homeostasis during skeletal muscle contraction,” Diabetes, vol. 53, no. 7, pp. 1643–1648, Jul. 2004, doi: 10.2337/DIABETES.53.7.1643.

[5] E. Hondares et al., “Thermogenic Activation Induces FGF21 Expression and Release in Brown Adipose Tissue,” J. Biol. Chem., vol. 286, no. 15, p. 12983, Apr. 2011, doi: 10.1074/JBC.M110.215889.

[6] G. I. Lancaster, K. Møller, B. Nielsen, N. H. Secher, M. A. Febbraio, and L. Nybo, “Exercise induces the release of heat shock protein 72 from the human brain in vivo,” Cell Stress Chaperones, vol. 9, no. 3, p. 276, Sep. 2004, doi: 10.1379/CSC-18R.1.

[7] P. Mera et al., “Osteocalcin signaling in myofibers is necessary and sufficient for optimum adaptation to exercise,” Cell Metab., vol. 23, no. 6, p. 1078, Jun. 2016, doi: 10.1016/J.CMET.2016.05.004.

[8] J. Hansen et al., “Exercise Induces a Marked Increase in Plasma Follistatin: Evidence That Follistatin Is a Contraction-Induced Hepatokine,” Endocrinology, vol. 152, no. 1, pp. 164–171, Jan. 2011, doi: 10.1210/EN.2010-0868.

[9] M. C. K. Severinsen and B. K. Pedersen, “Muscle–Organ Crosstalk: The Emerging Roles of Myokines,” Endocr. Rev., vol. 41, no. 4, pp. 594–609, Aug. 2020, doi: 10.1210/ENDREV/BNAA016.

[10] B. K. Pedersen, “The Physiology of Optimizing Health with a Focus on Exercise as Medicine,” Annual Review of Physiology, vol. 81. Annual Reviews Inc., pp. 607–627, Feb. 10, 2019, doi: 10.1146/annurev-physiol-020518-114339.

[11] B. K. Pedersen, “Muscles and their myokines,” J. Exp. Biol., vol. 214, no. 2, pp. 337 LP – 346, Jan. 2011, doi: 10.1242/jeb.048074.

[12] M. Yáñez-Mó et al., “Biological properties of extracellular vesicles and their physiological functions,” J. Extracell. vesicles, vol. 4, p. 27066, May 2015, doi: 10.3402/jev.v4.27066.

[13] D. Armstrong and D. E. Wildman, “Extracellular vesicles and the promise of continuous liquid biopsies,” J. Pathol. Transl. Med., vol. 52, no. 1, pp. 1–8, Jan. 2018, doi: 10.4132/JPTM.2017.05.21.

[14] K. Sidhom, P. O. Obi, and A. Saleem, “A Review of Exosomal Isolation Methods: Is Size Exclusion Chromatography the Best Option?,” Int. J. Mol. Sci. 2020, *Vol.* 21, *Page* 6466, vol. 21, no. 18, p. 6466, Sep. 2020, doi: 10.3390/IJMS21186466.

[15] J. A. Welsh et al., “Minimal information for studies of extracellular vesicles (MISEV2023): From basic to advanced approaches,” J. Extracell. vesicles, vol. 13, no. 2, Feb. 2024, doi: 10.1002/JEV2.12404.

[16] A. Safdar, A. Saleem, and M. A. Tarnopolsky, “The potential of endurance exercise-derived exosomes to treat metabolic diseases,” Nat. Rev. Endocrinol., vol. 12, no. 9, pp. 504–517, 2016, doi: 10.1038/nrendo.2016.76.

[17] M. Whitham et al., “Extracellular Vesicles Provide a Means for Tissue Crosstalk during Exercise,” Cell Metab., vol. 27, no. 1, pp. 237–251.e4, Jan. 2018, doi: 10.1016/J.CMET.2017.12.001/ATTACHMENT/CEDD5019-D644-47C6-8230-3188B7918EE0/MMC2.XLSX.

[18] C. Frühbeis, S. Helmig, S. Tug, and P. Simon & Eva-Maria Krämer-Albers, “Physical exercise induces rapid release of small extracellular vesicles into the circulation,” J. Extracell. Vesicles, 2015, doi: 10.3402/jev.v4.28239.

[19] Z. Hou et al., “Longterm Exercise-Derived Exosomal miR-342-5p: A Novel Exerkine for Cardioprotection,” Circ. Res., vol. 124, no. 9, pp. 1386–1400, Apr. 2019, doi: 10.1161/CIRCRESAHA.118.314635.

[20] C. Ma et al., “Moderate Exercise Enhances Endothelial Progenitor Cell Exosomes Release and Function,” Med. Sci. Sports Exerc., vol. 50, no. 10, pp. 2024–2032, Oct. 2018, doi: 10.1249/MSS.0000000000001672.

[21] P. Chaturvedi, A. Kalani, I. Medina, A. Familtseva, and S. C. Tyagi, “Cardiosome mediated regulation of MMP9 in diabetic heart: role of mir29b and mir455 in exercise,” J. Cell. Mol. Med., vol. 19, no. 9, pp. 2153–2161, Sep. 2015, doi: 10.1111/JCMM.12589.

[22] A. E. Rigamonti et al., “Effects of an acute bout of exercise on circulating extracellular vesicles: tissue-, sex-, and BMI-related differences,” Int. J. Obes. (Lond)., vol. 44, no. 5, pp. 1108–1118, May 2020, doi: 10.1038/S41366-019-0460-7.

[23] T. M. Pierdoná et al., “Extracellular vesicles as predictors of individual response to exercise training in youth living with obesity,” bioRxiv, p. 2020.11.20.390872, Nov. 2020, doi: 10.1101/2020.11.20.390872.

[24] J. A. C. Lovett, P. J. Durcan, and K. H. Myburgh, “Investigation of Circulating Extracellular Vesicle MicroRNA Following Two Consecutive Bouts of Muscle-Damaging Exercise,” Front. Physiol., vol. 9, no. AUG, Aug. 2018, doi: 10.3389/FPHYS.2018.01149.

[25] L. Sadovska et al., “Exercise-Induced Extracellular Vesicles Delay the Progression of Prostate Cancer,” Front. Mol. Biosci., vol. 8, p. 1346, Jan. 2022, doi: 10.3389/FMOLB.2021.784080.

[26] G. Annibalini et al., “Muscle and systemic molecular responses to a single flywheel based iso-inertial training session in resistance-trained men,” Front. Physiol., vol. 10, no. MAY, p. 554, 2019, doi: 10.3389/FPHYS.2019.00554.

[27] P. O. Obi, et al., “Chronic contractile activity induced skeletal muscle-derived extracellular vesicles increase mitochondrial biogenesis in recipient myocytes via transmembrane or peripheral membrane proteins,” bioRxiv, p. 2024.08.01.605916, Aug. 2024, doi: 10.1101/2024.08.01.605916.

[28] P. O. Obi, T. F. G. Souza, K. J. McManus, A. R. West, J. W. Gordon, and A. Saleem, “The pro-apoptotic effect of chronic contractile activity-induced extracellular vesicles on Lewis Lung Carcinoma cells,” bioRxiv, p. 2024.08.29.610232, Aug. 2024, doi: 10.1101/2024.08.29.610232.

[29] B. MM, “A rapid and sensitive method for the quantitation of microgram quantities of protein utilizing the principle of protein-dye binding,” Anal. Biochem., vol. 72, no. 1–2, pp. 248–254, May 1976, doi: 10.1006/ABIO.1976.9999.

[30] M. Pathan et al., “FunRich: An open access standalone functional enrichment and interaction network analysis tool,” Proteomics, vol. 15, no. 15, pp. 2597–2601, Aug. 2015, doi: 10.1002/PMIC.201400515.

[31] S. Keerthikumar et al., “ExoCarta: A web-based compendium of exosomal cargo,” J. Mol. Biol., vol. 428, no. 4, p. 688, Feb. 2016, doi: 10.1016/J.JMB.2015.09.019.

[32] M. Pathan et al., “Vesiclepedia 2019: a compendium of RNA, proteins, lipids and metabolites in extracellular vesicles,” Nucleic Acids Res., vol. 47, no. Database issue, p. D516, Jan. 2019, doi: 10.1093/NAR/GKY1029.

[33] D. Szklarczyk et al., “The STRING database in 2021: customizable protein–protein networks, and functional characterization of user-uploaded gene/measurement sets,” Nucleic Acids Res., vol. 49, no. D1, p. D605, Jan. 2021, doi: 10.1093/NAR/GKAA1074.

[34] A. Krämer, J. Green, J. Pollard, and S. Tugendreich, “Causal analysis approaches in Ingenuity Pathway Analysis,” Bioinformatics, vol. 30, no. 4, pp. 523–530, Feb. 2014, doi: 10.1093/BIOINFORMATICS/BTT703.

[35] A. Forterre et al., “Proteomic Analysis of C2C12 Myoblast and Myotube Exosome-Like Vesicles: A New Paradigm for Myoblast-Myotube Cross Talk?,” PLoS One, vol. 9, no. 1, Jan. 2014, doi: 10.1371/JOURNAL.PONE.0084153.

[36] S. Watanabe et al., “Skeletal muscle releases extracellular vesicles with distinct protein and microRNA signatures that function in the muscle microenvironment,” PNAS Nexus, vol. 1, no. 4, Oct. 2022, doi: 10.1093/PNASNEXUS/PGAC173.

[37] A. S. Deshmukh, M. Murgia, N. Nagaraj, J. T. Treebak, J. Cox, and M. Mann, “Deep Proteomics of Mouse Skeletal Muscle Enables Quantitation of Protein Isoforms, Metabolic Pathways, and Transcription Factors,” Mol. Cell. Proteomics, vol. 14, no. 4, p. 841, Apr. 2015, doi: 10.1074/MCP.M114.044222.

[38] M. M. Savitski et al., “Measuring and managing ratio compression for accurate iTRAQ/TMT quantification,” J. Proteome Res., vol. 12, no. 8, pp. 3586–3598, Aug. 2013, doi: 10.1021/PR400098R/SUPPL_FILE/PR400098R_SI_001.PDF.

[39] M. C. Chong et al., “Acute exercise-induced release of innate immune proteins via small extracellular vesicles changes with aerobic fitness and age,” Acta Physiol., vol. 240, no. 3, p. e14095, Mar. 2024, doi: 10.1111/APHA.14095.

[40] A. Brahmer et al., “Platelets, endothelial cells and leukocytes contribute to the exercise-triggered release of extracellular vesicles into the circulation,” J. Extracell. Vesicles, vol. 8, p. 1615820, Dec. 2019, doi: 10.1080/20013078.2019.1615820.

[41] M. Guescini et al., “Muscle Releases Alpha-Sarcoglycan Positive Extracellular Vesicles Carrying miRNAs in the Bloodstream,” PLoS One, vol. 10, no. 5, May 2015, doi: 10.1371/JOURNAL.PONE.0125094.

[42] A. L. Estrada et al., “Deconstructing Organs: Single-Cell Analyses, Decellularized Organs, Organoids, and Organ-on-a-Chip Models: Extracellular vesicle secretion is tissue-dependent ex vivo and skeletal muscle myofiber extracellular vesicles reach the circulation in vivo,” Am. J. Physiol. -Cell Physiol., vol. 322, no. 2, p. C246, Feb. 2022, doi: 10.1152/AJPCELL.00580.2020.

[43] W. Aoi et al., “A novel myokine, secreted protein acidic and rich in cysteine (SPARC), suppresses colon tumorigenesis via regular exercise,” Gut, vol. 62, no. 6, pp. 882–889, Jun. 2013, doi: 10.1136/GUTJNL-2011-300776.

[44] C. A. Henderson, C. G. Gomez, S. M. Novak, L. Mi-Mi, and C. C. Gregorio, “Overview of the Muscle Cytoskeleton,” Compr. Physiol., vol. 7, no. 3, p. 891, Jun. 2017, doi: 10.1002/CPHY.C160033.

[45] I. Y. Kuo and B. E. Ehrlich, “Signaling in Muscle Contraction,” Cold Spring Harb. Perspect. Biol., vol. 7, no. 2, 2015, doi: 10.1101/CSHPERSPECT.A006023.

[46] L. S. Holliday, L. P. de Faria, and W. J. Rody, “Actin and Actin-Associated Proteins in Extracellular Vesicles Shed by Osteoclasts,” Int. J. Mol. Sci., vol. 21, no. 1, Jan. 2020, doi: 10.3390/IJMS21010158.

[47] P. Fuentes et al., “ITGB3-mediated uptake of small extracellular vesicles facilitates intercellular communication in breast cancer cells,” Nat. Commun. 2020 111, vol. 11, no. 1, pp. 1–15, Aug. 2020, doi: 10.1038/s41467-020-18081-9.

[48] D. Shimura and R. M. Shaw, “GJA1-20k and Mitochondrial Dynamics,” Front. Physiol., vol. 13, Mar. 2022, doi: 10.3389/FPHYS.2022.867358.

[49] T. A. Read et al., “The actin binding protein profilin 1 localizes inside mitochondria and is critical for their function,” EMBO Rep., Aug. 2024, doi: 10.1038/S44319-024-00209-3/SUPPL_FILE/44319_2024_209_MOESM8_ESM.PDF.

[50] G. Nowak and J. Megyesi, “Protein kinase Cα mediates recovery of renal and mitochondrial functions following acute injury,” FEBS J., vol. 287, no. 9, pp. 1830–1849, May 2020, doi: 10.1111/FEBS.15110.

[51] S. Riis, J. B. Murray, and R. O’Connor, “IGF-1 Signalling Regulates Mitochondria Dynamics and Turnover through a Conserved GSK-3β–Nrf2–BNIP3 Pathway,” Cells, vol. 9, no. 1, Jan. 2020, doi: 10.3390/CELLS9010147.

[52] A. Bhattacharjee et al., “The Activity of Menkes Disease Protein ATP7A Is Essential for Redox Balance in Mitochondria,” J. Biol. Chem., vol. 291, no. 32, p. 16644, Aug. 2016, doi: 10.1074/JBC.M116.727248.

[53] D. De Rasmo, A. Signorile, E. Roca, and S. Papa, “cAMP response element-binding protein (CREB) is imported into mitochondria and promotes protein synthesis,” FEBS J., vol. 276, no. 16, pp. 4325– 4333, Aug. 2009, doi: 10.1111/J.1742-4658.2009.07133.X.

[54] A. D. Cherry and C. A. Piantadosi, “Regulation of mitochondrial biogenesis and its intersection with inflammatory responses,” Antioxid. Redox Signal., vol. 22, no. 12, pp. 965–976, Apr. 2015, doi: 10.1089/ARS.2014.6200.

[55] Y. W. Zhang, I. Morita, M. Ikeda, K. W. Ma, and S. Murota, “Connexin43 suppresses proliferation of osteosarcoma U2OS cells through post-transcriptional regulation of p27,” Oncogene, vol. 20, no. 31, pp. 4138–4149, Jul. 2001, doi: 10.1038/SJ.ONC.1204563.

[56] Y. Wang, Y. Wang, R. Wan, C. Hu, and Y. Lu, “Profilin 1 Protein and Its Implications for Cancers,” Oncology (Williston Park)., vol. 35, no. 7, pp. 402–409, Jul. 2021, doi: 10.46883/ONC.2021.3507.0402.

[57] Y. J. Liu and C. Wang, “A review of the regulatory mechanisms of extracellular vesicles-mediated intercellular communication,” Cell Commun. Signal., vol. 21, no. 1, p. 77, Dec. 2023, doi: 10.1186/S12964-023-01103-6.

