## supplementary figures for "Identification of novel myokines and putative protein targets that mediate functional adaptations in response to chronic contractile activity induced skeletal muscle-extracellular vesicle treatment"

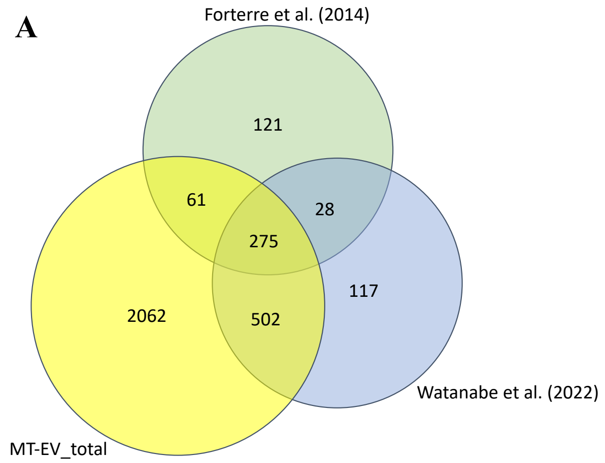


**Figure S1. Comparison of EV proteome in present study vs. previous studies.** Venn diagrams showing the total proteins (MT-EV_total) compared with proteins from previous studies. Comparison was done between **(A)** MT-EV_total vs. myotube-EV proteome (from Forterre et al., 2014 [35], and Watanabe et al., 2022 [36]), and **(B)** MT-EV_total vs. significantly regulated EV proteins following acute exercise (from Whitham et al., 2018 [17]).

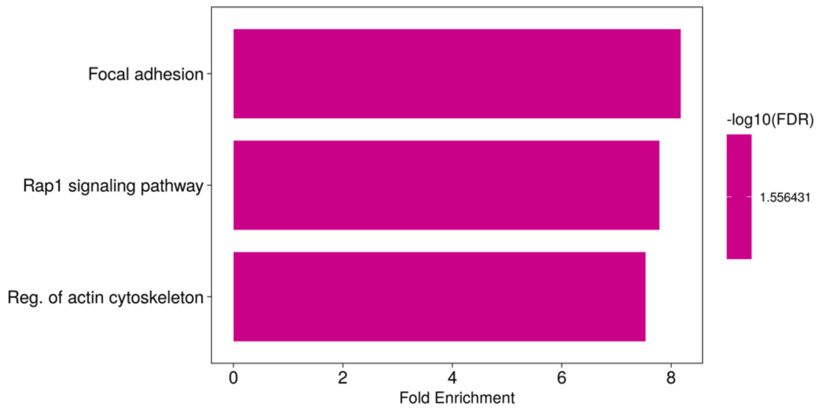


**Figure S2. KEGG analysis of differentially expressed proteins.** The top 3 canonical pathways associated with the DEPs are identified by KEGG analysis with −log10 (FDR) values and fold enrichment indicated.
